# A novel immunogene therapy to cancer with high tumor selectivity and safety

**DOI:** 10.1101/2025.03.14.643237

**Authors:** Yile Wang, Ziyan Kong, Yunqi Zhao, Bing Pei, Jian Sun, Xueyuan Mao, Weida Gong, Ying Chen, Rong Yin, Meng Cao, Jinke Wang

**Affiliations:** School of Biological Science and Medical Engineering, Southeast University, Nanjing 210096, China; Suqian Clinical College, Xuzhou Medical University, Xuzhou, China; Suqian First People’s Hospital, Nanjing Medical University, Nanjing, China; Yixing People’s Hospital, Yixing, China; School of Medical Technology, Xuzhou Medical University, Xuzhou 221004, China; Jiangsu Cancer Hospital, Nanjing Medical University, Nanjing 21009, China; Division of Breast Surgery, Department of General Surgery, Nanjing Drum Tower Hospital, the Affiliated Hospital of Medical School, Nanjing University, Nanjing, 210008, China

**Author notes:** To whom correspondence should be addressed: Jinke Wang, Sipailou 2, School of Biological Science and Medical Engineering, Southeast University, Nanjing 210096, China.

**Keywords:** Immunogene therapy, immunotherapy, immune checkpoints, cytokine, adeno-associated virus

## Abstract

Cancer immunotherapy has made significant advancements over the past few decades, with immune checkpoint and cytokine-based drugs being successfully implemented in clinical settings. Nonetheless, the effective and safe clinical application of these therapies is hindered by critical issues, such as severe toxicity to healthy tissues due to on-target off-tumor effects. In this study, we have developed a novel immunogene therapy characterized by high tumor selectivity and safety in vivo, effectively mitigating the off-tumor effects associated with current antibody-based immune checkpoint therapies. We engineered a gene expression vector that is specifically activated by NF-κB activity to co-express artificial microRNAs targeting two key immune checkpoints (PD-L1 and CD47) and cytokine IL-15. This vector is capable of selectively and effectively down regulating the expression of PDL1 and CD47 while over expressing IL-15 just exclusively in cancer cells, both in vitro and in vivo. Through this mechanism, both adaptive and innate immune responses can be simultaneously activated and enhanced via the transfection of this vector. The in vivo administration of this vector via recombinant adeno-associated virus (AAV) demonstrated significant antitumor activity, high tumor selectivity, and safety in murine models. Consequently, this vector may offer a potential more effective and safer alternative to the current immune checkpoint inhibitors in future clinical applications.

## Introduction

Cancer is a severe, and life-threatening condition, characterized by high incidence and mortality rates [1, 2], and is recognized for its complexity [3–5]. In recent decades, substantial efforts have been directed towards combating cancer through the development of various medical and pharmaceutical technologies, including surgical interventions, chemotherapy, radiotherapy, targeted therapy, and immunotherapy [6]. Notably, immunotherapy has garnered increasing attention in both biomedical research and clinical practice. The development of cancer is often attributed to a failure of immune surveillance, which allows cancer cells to evade immune detection. Immunotherapies are designed to harness the immune system’s capacity to identify and eliminate cancer cells [7]. These therapeutic strategies encompass oncolytic virus therapy [8], tumor vaccine therapy [9], monoclonal antibody therapy [10], antibody-drug conjugates (ADCs) [11], cytokine therapy [12], immune checkpoint inhibitor (ICI) therapy [13], and adoptive cell therapy (ACT) [14–16]. ICIs function by targeting tumor-induced immune tolerance, thereby reactivating immune cells to combat cancer cells through the utilization of antibodies that inhibit immune checkpoints (ICs), such as cytotoxic T-lymphocyte-associated antigen 4 (CTLA-4) [17], programmed cell death protein 1 (PD-1) and programmed cell death-ligand 1 (PD-L1) [18, 19]. Nevertheless, each type of immunotherapy exhibits distinct advantages and limitations in clinical settings, primarily due to the substantial tumor heterogeneity observed both within individual patients and across different patients.

The PD-1 and PD-L1 molecules constitute a pair of co-inhibitory entities crucial for immune regulation. PD-1 is present on the membrane surfaces of various immune cells, including T and B lymphocytes, natural killer (NK) cells, macrophages, and dendritic cells (DCs) [18, 20]. In addition to its expression on immune cells, PD-L1 is ubiquitously expressed on a variety of healthy tissue cells, such as vascular endothelial cells, keratinocytes, pancreatic islet cells, astrocytes, placenta syncytiotrophoblast cells, and corneal epithelial and endothelial cells [21]. Consequently, the PD-1/PD-L1 axis is regarded as a pivotal immune checkpoint (IC) that protects normal cells from immune cell attacks, thereby immune tolerance. However, PD-L1 is also expressed on cancer cells [22], facilitating immune evasion and posing a significant obstacle to cancer treatment [23]. To counteract this, PD-1/PD-L1 antibodies have been developed and successfully implemented in clinical cancer treatment to block immune escape [24]. Nevertheless, the clinical application of these antibodies as ICIs faces challenges, including limited responses (approximately 30% in solid tumors) [25] and severe immune-related adverse events (irAEs), primarily resulting from on-target off-tumor effects [26–28]. Consequently, there remains a significant demand for the development of more effective and safer immunotherapies targeting PD-1/PD-L1.

In addition to ICs that primarily regulate T cell-mediated adaptive immunity, such as PD-1/PD-L1, there is a growing interest in ICs that predominantly modulate innate immunity. An illustrative example is CD47, a cell-surface immunomodulatory molecule that ubiquitously expressed across various cell types, which provides protection against immune cell attacks, including those form macrophages and natural killer (NK) cells. CD47 is commonly described as a “don’t eat me” signal, as it prevents phagocytosis by binding to signal regulatory protein α (SIRPα) on the surface of macrophages [29, 30]. Tumor cells exploit this mechanism by overexpressing CD47 on the their surfaces to evade phagocytosis [31, 32]. As a result, CD47 has emerged as a promising target for cancer immunotherapy. Nevertheless, the development of inhibitors, such as antibodies against CD47, is significantly hindered severe on-target off-tumor side effects. Notably, CD47 antibodies have been associated with hematotoxicity to erythrocytes, potentially leading to anemia and thrombocytopenia [33]. Therefore, there is an urgent need for the development of safer immunotherapies targeting CD47.

Effective immunotherapy is contingent upon the robust activity of immune cells, which in turn is dependent on chemokines and cytokines with immune-stimulating properties. Consequently, cytokine-based immunotherapy has been investigated for a longer duration compared to ICIs. For instance, large-dose Interleukin-12 (IL-2) has been shown to significantly reduce tumor regression in patients with metastatic tumors [34]. Interleukin-15 (IL-15) is another pivotal cytokine produced by various cells, including DCs, macrophages, and fibroblasts [35]. IL-15 facilitates the growth and proliferation of T and NK cells, impacts memory cells, enhances cell-mediated immune responses, and augments the migratory capability of immune cells in antitumor responses [36, 37]. Consequently, IL-15 is regarded as a promising immunotherapeutic agent that can enhance the antitumor efficacy of ICIs [38]. However, systematic administration of IL-15 at effective doses often leads to excessive immune activation and subsequent systematic toxicity [39, 40], necessitating localized administrated to the tumor site to mitigate toxicity [41]. Therefore, the development of safer IL-15-based immunotherapies remains an unmet need in the filed biomedicine.

In prior research, we demonstrated a cancer cell-specific gene expression technology predicated on NF-κB activity within cancer cells [42]. Utilizing this technology, a cancer ferroptosis therapy was developed [43, 44], underscoring the high tumor selectivity and safety of the NF-κB-activatable therapeutic gene expression system. Building upon this technology, we have developed an innovative immunogene therapy aimed at addressing the clinical challenges associated with current ICIs-based immunotherapies. Three recombinant adeno-associated viruses (rAAVs), including rAAV-miPDL1, rAAV-miCD47, and rAAV-mIL15, were engineered to enhance antitumor activity. Specifically, rAAV-miPDL1 was designed to provoke the antitumor activity of cytotoxic T lymphocytes (CTLs) by selectively knocking down PD-L1 in tumor cells; rAAV-miCD47 was developed to stimulate the antitumor activities of macrophages and NK cells by specifically targeting CD47 in tumor cells; and rAAV-mIL15 was constructed to augment the antitumor activities of immune cells through the overexpression of IL-15 in tumor cells. Ultimately, a combinatorial rAAV, termed rAAV-miPDL1-miCD47-IL15, was developed to co-express PDL1- and CD47-targeting microRNAs (miRs) along with the IL-15 cytokine, thereby achieving a synergetic therapeutic effect. All rAAV constructs demonstrated significant antitumor efficacy, high tumor selectivity and safety in murine models.

## Materials and methods

### 1. Vector Construction

The sequences of murine PD-L1, CD47, and IL-15 were obtained from the NCBI database. miRNAs targeting murine and human PD-L1 and CD47 were designed using the BLOCK-iT™ RNAi Designer (https://rnaidesigner.thermofisher.com/rnaiexpress/sort.do) (Additional file 1: Table S1). Utilizing the specific promoter vector pDMP-miR developed in our laboratory, we constructed the vectors pDMP-miPDL1, pDMP-miCD47, and pDMP-IL15. Notably, during the construction of pDMP-IL15, an enhanced expression element, WPRE, was incorporated. The gene expression cassettes of these vectors were subsequently digested with restriction endonucleases and incorporated into a recombinant adeno-associated viral vector (pAAV-MCS) to generate pAAV-miPDL1, pAAV-miCD47, and pAAV-IL15. Co-expression vectors, pAAV-miPDL1-miCD47 and pAAV-miPDL1-miCD47-IL15, were constructed via homologous recombination cloning. Negative control vectors, designated as pAAV-miNT and pAAV-MCS-miNT, were constructed similarly, where miNT encodes a miR targeting no transcripts (NT) and MCS encodes no transcripts. These vectors were developed for both human and murine genes and were utilized in the treatment of human and murine cells, as well as murine models.

### 2. Cells Culture

The cell lines utilized in this study comprised HEK-293 (human embryonic kidney cells), CT26 (murine colon cancer cells), NIH-3T3 (murine embryonic fibroblasts), CTLL2 (murine T-lymphocytes), RAW264.7 (murine mononuclear macrophages), MRC-5 (human embryonic lung fibroblasts), A549 (human non-small cell lung cancer cells), HepG2 (human hepatocellular carcinoma cells), HCT116 (human colon cancer cells), Jurkat (human T cell leukemia cells), and THP-1 (human monocytes). These cell lines were procured from the Cell Resource Center of the Shanghai Institutes for Biological Sciences, affiliated with the Chinese Academy of Sciences. HEK-293T, NIH-3T3, RAW264.7, MRC-5, A549, HepG2, and HCT116 cells were cultured in Dulbecco’s Modified Eagle Medium (DMEM) supplied by Gibco. CT26 and Jurkat cells were cultured in Roswell Park Memorial Institute (RPMI) 1640 medium, also from Gibco. THP-1 and CTLL2 cells were maintained in specialized media provided by Servicebio. All culture media were supplemented with 10% fetal bovine serum (HyClone), along with 100 units/mL penicillin and 100 μg/mL streptomycin (Thermo Fisher). The cells were incubated at 37 °C in a humidified atmosphere containing 5% CO2.

### 3. Cell co-culture

In the co-culture experiment involving Jurkat and HCT116 cells, HCT116 cells were initially transfected with the plasmids pAAV-miPDL1 or pAAV-miNT for a duration of 48 hours. Subsequently, these cells were seeded into 96-well plates at a density of 10,000 cells per well and cultured for an additional 24 hours. Concurrently, Jurkat cells were activated using PMA at a concentration of 50 ng/mL and PHA at 2 μg/mL for 12 hours. Following activation, the transfected HCT116 cells were co-cultured with the activated Jurkat cells at a ratio of 4:1 (Jurkat:HCT116) for 36 hours. Post co-culture, Jurkat cells were harvested, and their mRNA expression levels of IL-2 and IFN-γ were quantified using qPCR. Meanwhile, the HCT116 cells were replenished with fresh culture medium, and their viability was assessed using the CCK-8 assay.

In a separate co-culture experiment involving THP-1 and A549/HCT116 cells, A549/HCT116 cells were transfected with either pAAV-miNT or pAAV-miCD47 plasmids and cultured for 72 hours. THP-1 cells were induced to differentiate with PMA at 100 ng/mL for 48 hours, followed by polarization through the addition of LPS at 100 ng/mL in the presence of PMA for 24 hours. The polarized THP-1 cells were then co-cultured with the transfected A549/HCT116 cells at a ratio of 10:1 (THP-1:HCT116) for 48 hours. After co-culture, the cells were refreshed with new culture medium, and cell viability was determined using the CCK-8 reagent.

The co-culture of RAW264.7 and CT26 cells involved the transfection of CT26 cells with pAAV-NT or pAAV-miCD47 for a duration of 72 hours. Concurrently, RAW264.7 cells were induced to differentiate using PMA at a concentration of 50 ng/mL for 48 hours. Subsequently, the differentiated RAW264.7 cells were co-cultured with the transfected CT26 cells for 20 hours at a ratio of 10:1 (RAW264.7:CT26). Following this period, the culture medium was refreshed, and cell viability was assessed using the CCK-8 assay.

In a parallel experiment, the co-culture of Jurkat and HCT116 cells was conducted as follows: HCT116 cells were transfected with the plasmid pAAV-MCS or pAAV-hIL15 for 48 hours, after which they were seeded into 96-well plates at a density of 10,000 cells per well and cultured for an additional 24 hours. Jurkat cells were activated with PMA at 50 ng/mL and PHA at 2 μg/mL for 12 hours. The activated Jurkat cells were then co-cultured with the transfected HCT116 cells at a ratio of 5:1 (Jurkat:HCT116) for 36 hours. The viability of the co-cultured cells was subsequently determined using the CCK-8 assay.

### 4. T cell activation in vitro

In the CTLL-2 activation assay, CTLL-2 cells, which are a transformed cell line inherently independent of activation, exhibit growth dependence on IL-2 at a concentration of 100 ng/mL. Mouse CT26 cells were transfected with the plasmids pAAV-MCS and pAAV-mIL15, respectively, and the supernatant was harvested after 72 hours. This supernatant was subsequently mixed with CTLL-2-specific medium in a 1:1 ratio to culture the CTLL-2 cells. The viability of the CTLL-2 cells was assessed using the CCK-8 reagent after a 60-hour incubation period.

### 5. Virus Preparation

HEK-293T cells were seeded into 75 cm² flasks at a density of 5 × 10⁶ cells per flask and cultured until reaching 80% confluence. Subsequently, the cells were co-transfected with two helper plasmids (pHelper and pAAV-RC; Stratagene) and one pAAV plasmid using Lipofectamine 2000, following the manufacturer’s instructions (Thermo Fisher Scientific). After 72 hours, both the cells and media were harvested and subjected to three freeze-thaw cycles, initially freezing at −80°C and then thawing in a 37°C water bath. Chloroform was added to the lysate at a ratio of 1:10 (v/v), and the mixture was vigorously agitated at 37°C for 1 hour. Sodium chloride was added to achieve a final concentration of 1 M, and the solution was mixed until the NaCl was fully dissolved, followed by centrifugation at 15,000 rpm at 4°C for 15 minutes to collect the supernatant. Polyethylene glycol 8000 (PEG8000) was then added to the supernatant to a final concentration of 10% (w/v), and the solution was mixed until dissolved and incubated on ice for 1 hour. The mixture was centrifuged again at 15,000 rpm at 4°C for 15 minutes. The supernatant was discarded, and the resulting pellet was resuspended in PBS. DNase and RNase were added to the mixture at a final concentration of 1 μg/mL, and the solution was incubated at room temperature for 30 minutes. The mixture was extracted with chloroform at a 1:1 volume ratio, and the aqueous layer containing the purified virus was transferred to a new tube. The titers of the adeno-associated viruses (AAVs) were determined using quantitative PCR (qPCR) with the primers AAV-F/R, as detailed in Additional File 1: Table S2. The quantified viruses were aliquoted and stored at -80°C. The resulting viruses were designated as rAAV-miNT, rAAV-MCS, rAAV-MCS-NT, rAAV-miCD47, rAAV-miPDL1, rAAV-IL15, rAAV-miPDL1-miCD47, and rAAV-miPDL1-miCD47-IL15, respectively.

### 6. Quantitative PCR

Total RNA was extracted from cells or mouse tissues utilizing TRIzol™ (Invitrogen) in accordance with the manufacturer’s instructions. Complementary DNA (cDNA) synthesis was conducted using the FastKing RT kit (Tiangen). Genomic DNA (gDNA) was isolated from various mouse tissues employing the TIANamp Genomic DNA Kit (TIANGEN). Gene amplification from both cDNA and gDNA was carried out via quantitative PCR (qPCR) using the Hieff qPCR SYBR Green Master Mix (Yeasen). Each treatment condition was assessed in triplicate on an ABI Step One Plus system (Applied Biosystems). Relative mRNA transcript levels were normalized to the GADPH internal reference and calculated as relative quantity (RQ) using the formula: RQ = 2^‾ΔΔCt^. Similarly, viral DNA abundance was normalized to the GADPH internal reference and calculated using the formula: RQ = 2^‾–ΔCt^. Specificity of all qPCR primers was confirmed through melting curve analysis, with primer sequences detailed in Additional file 1: Table S2. All experimental procedures were conducted in triplicate.

### 7. ELISA

The plasmid pAAV-DMP-mIL15-WPRE was transfected into CT26 cells, and the supernatant was collected for analysis via enzyme-linked immunosorbent assay (ELISA) after 72 hours. The experiment employed an Elabscience Mouse IL-15 ELISA Kit (E-EL-M0728), which utilizes a double antibody sandwich ELISA technique. In this method, the enzyme labeling plate is coated with an anti-Mouse IL-15 antibody, which captures the mouse IL-15 present in the samples. Subsequently, a biotinylated anti-mouse IL-15 antibody and a horseradish peroxidase-labeled affinity element are added sequentially. The biotinylated antibody binds to the mouse IL-15 already attached to the coated antibody, while the biotin and affinity element form a specific immune complex. The chromogenic substrate, tetramethylbenzidine (TMB), is catalyzed by horseradish peroxidase to produce a blue color, which turns yellow upon the addition of a stop solution. The optical density (OD) was measured at 450 nm using a spectrophotometer. A direct correlation was observed between the IL-15 concentration and the absorbance at 450 nm (A450), allowing for the determination of IL-15 concentration in the samples by constructing a standard curve with known standards.

### 8. AO & EB staining

Jurkat and HCT116 cells were co-cultured, with the Jurkat cells maintained in suspension, allowing for their removal via fluid exchange, thereby isolating the adherent HCT116 cells. Subsequently, the HCT116 cells were stained using acridine orange and ethidium bromide (AO&EB, Solarbio) in accordance with the manufacturer’s protocol. Fluorescence microscopy (IX51, Olympus) was employed to visualize and quantify the live and dead cell populations.

### 9. Animal Treatments

Four-week-old female BALB/c mice were procured from Changzhou Cavens Laboratory Animal Co. Ltd (China). All animal experiments conducted in this study adhered to the guidelines and ethical standards set by the Animal Care and Use Committee of Southeast University (Nanjing, China). Tumor volumes were assessed using Vernier calipers, applying the formula V = (ab^2^)/2, where ’a’ represents the longest diameter and ’b’ the shortest diameter. Euthanasia was performed on mice when tumors reached a volume of 2000 mm^3^ or when there was a 20% loss in body weight. Various tissues, including the heart, liver, spleen, lung, kidney, and tumor, were harvested for subsequent analyses. CT26 xenografts were established in BALB/c mice through the subcutaneous injection of 1 × 10^6^ CT26 cells into the inner thighs. The mice were maintained for 10 days to allow for tumor development, achieving a tumor volume exceeding 150 mm^3^. Mice were then randomly assigned to groups and received intravenous injections every two days, totaling three administrations, with different reagents, including PBS and various recombinant adeno-associated viruses (rAAVs). Each injection consisted of 100 μl of either PBS or rAAV, with the rAAV injections containing 5 × 10^10^ viral genomes (vg) per mouse. Tumor dimensions and body weight of the mice were assessed daily. On the seventh day following viral injection (corresponding to the seventeenth day after tumor inoculation), the mice were euthanized and photographed. The rAAVs utilized in the animal experiments included rAAV-miNT, rAAV-MCS, rAAV-MCS-NT, rAAV-miPDL1, rAAV-miCD47, rAAV-mIL15, rAAV-miPDL1-miCD47, and rAAV-miPDL1-miCD47-mIL15.

### 10. Hematoxylin and eosin (H&E) staining

Individual tissues, including the heart, liver, spleen, lungs, kidneys, and tumors, were harvested from mice and initially fixed in a 4% paraformaldehyde solution to preserve their morphology and structural integrity. Following fixation, the tissues underwent a dehydration process using a graded series of alcohol concentrations. Subsequently, the tissues were embedded in paraffin wax and sectioned into thin slices using a microtome. The paraffin was then removed with xylene, and the sections were rehydrated through a series of alcohol solutions. For histological examination, the nuclei were stained with hematoxylin, imparting a blue-violet hue, while the cytoplasm was counterstained with eosin, resulting in a pink coloration. The prepared slides were subsequently imaged using an Olympus IX51 microscope.

### 11. CCK-8 assay

CT26 cells were transfected with the plasmids pAAV-MCS and pAAV-DMP-mIL15-WPRE, respectively, and the supernatant was harvested after 72 hours. This supernatant was then mixed in a 1:1 ratio with CTLL2-specific medium, and CTLL2 cell viability was assessed using the CCK-8 reagent (Vazyme) after 60 hours. The experimental procedure was as follows: cells were seeded into 96-well plates at a density of 10,000 cells per well and incubated for 24 hours. Subsequently, 10 μL of CCK-8 solution was added to each well, followed by a 4-hour incubation at 37°C. The cells were gently mixed, and the absorbance at 450 nm was measured using a microplate reader (BioTek). Cell viability was then calculated using the formula: % viability = [(As‒Ab) / (Ac‒Ab)] × 100, where As represents the sample optical density (OD), Ac the control OD, and Ab the blank OD.

### 12. Blood and biochemical detection

Mice were euthanized, and blood was collected via cardiac puncture through the chest cavity using a sterile syringe. Whole blood samples were stored and transported at 4 °C with an anticoagulant. The following parameters were analyzed: red blood cells (RBC), white blood cells (WBC), blood platelets (PLT), and hemoglobin (HGB) using a Myriad Veterinary Automated Blood Cell Analyzer (Servicebio). For serum processing, blood samples were centrifuged at 4 °C at 3000 rpm for 15 minutes within 30 minutes of collection. The supernatant was then collected and stored in an 80 °C storage box for transport, ensuring that repeated freezing and thawing were avoided. Serum samples were analyzed for the following biochemical parameters: alanine aminotransferase (ALT), aspartate aminotransferase (AST), alkaline phosphatase (ALP), blood urea nitrogen (BUN), creatinine (Cr), and uric acid (UA) using a Fully Automated Biochemistry Instrument (Servicebio).

### 13. Statistical Analysis

All data are expressed as mean values ± standard deviation (S.D.). Statistical analyses and graphical representations were conducted using GraphPad Prism version 10.0 software. Statistical differences between two groups were assessed using a two-tailed Student’s t-test. For comparisons involving three or more groups, one-way or two-way analysis of variance (ANOVA) with Tukey’s post hoc correction for multiple comparisons was employed. The Kaplan–Meier method was utilized to evaluate differences in animal survival, with P values calculated using the log-rank test. A significance threshold of P < 0.05 was applied to determine statistical significance.

## Results

### 1. PD-L1 knockdown to treat mouse colon cancer

To investigate the therapeutic potential of PD-L1 knockdown in mouse models of colon cancer, we focused on the PD-1/PD-L1 axis, which serves as an inhibitory signal for T cell activation. Inhibition of this pathway can enhance adaptive antitumor immunity. Tumor cells often exhibit high levels of PD-L1 expression, facilitating immune evasion. To counteract this mechanism and promote T cell-mediated cytotoxicity, we engineered miPDL1 constructs to specifically target and reduce PD-L1 expression in tumor cells (Fig. 1a). Three microRNAs (miRs) were designed to target murine PD-L1: mus-miPDL1a, mus-miPDL1b, and mus-miPDL1c. Expression vectors, namely pDMP-miPDL1a/b/c, were developed for transfection into the mouse colon cancer cell line CT26 and mouse embryonic fibroblast cell line 3T3. The results demonstrated that all miPDL1 constructs effectively reduced PD-L1 expression in CT26 cancer cells, without affecting PD-L1 levels in the normal 3T3 cells (Fig. 1b). This indicates the specificity and efficacy of the miPDL1 constructs in selectively targeting PD-L1 in cancer cells. Among the constructs, mus-miPDL1b exhibited the highest knockdown efficiency. Consequently, it was utilized to create the vector pAAV-miPDL1, which was employed to package rAAV-miPDL1 for in vivo treatment. Mice bearing CT26 tumors were subsequently treated with either rAAV-miNT or rAAV-miPDL1 (Fig. 1c). The findings demonstrate that the rAAV-miPDL1 treatment significantly suppresses tumor growth compared to rAAV-miNT (Fig. 1d). The therapeutic efficacy of rAAV-miPDL1 is further corroborated by tumor imaging (Fig. 1e), as well as measurements of tumor size and weight (Fig. 1f), and histological analysis via H&E staining of tumor tissues (Fig. 1g). Additionally, the therapeutic impact of rAAV-miPDL1 is evidenced by a significant reduction in spleen size and weight (Figs. 1e and 1f). Preliminary assessments of the treatment’s biosafety are suggested by its minimal impact on the body weight of mice (Additional file 1: Fig. S1a). This biosafety is largely attributed to the selective knockdown of PD-L1 in tumor tissues, without affecting other healthy tissues (Fig. 1h), even though rAAV is distributed across all tissues (Additional file 1: Fig. S1b). Furthermore, the treatment’s biosafety is supported by its negligible effects on blood biochemical markers (WBC, RBC, PLT, HGB) (Additional file 1: Fig. S1c), liver function markers (ALT, AST, ALP) (Additional file 1: Fig. S1d), kidney function markers (BUN, Cr, UA) (Additional file 1: Fig. S1d), and the structural integrity of healthy tissues (Additional file 1: Fig. S1e). Collectively, these data underscore the antitumor efficacy of the rAAV-miPDL1 treatment and its favorable safety profile.

**Fig. 1.**
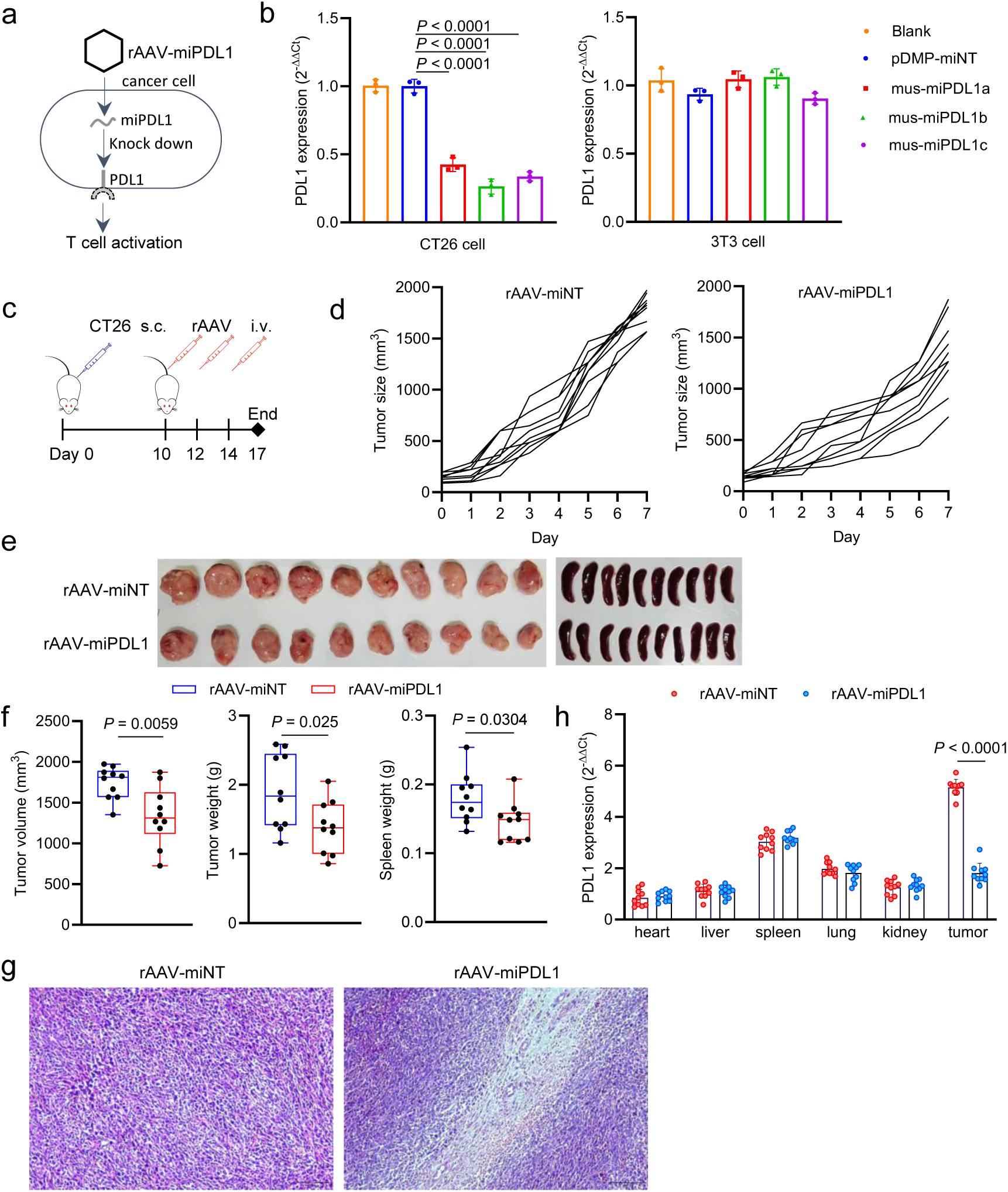
Treatment of CT26-xenografted mice using rAAV-miCD47. **a** Schematic representation of knocking down CD47 expression with rAAV-miCD47. **b** CD47 expression in CT26 and 3T3 cells following various transfections, as measured by qPCR (n=3 wells). **c** Killing activity of RAW264.7 cells against CT26 cells detected by CCK-8 reagent (n=3 wells). **d** Schematic overview of animal treatment protocol for CT26 xenograft mice, including subcutaneous (s.c.) and intravenous (i.v.) injections. **e** Tumor growth curve (n = 6 mice). **f** Imaging of the tumor and spleen. **g** Tumor volume, tumor weight, and spleen weight (n=6 mice). **h** Representative HE-staining tissue section of tumors. **i** Expression of CD47 levels in various tissues, as detected by qPCR (n=6 mice).

### 2. CD47 knockdown to treat mouse colon cancer

In addition to adaptive immunity, innate immunity plays a crucial role in the regulation of tumorigenesis and tumor progression, with macrophages and NK cells being pivotal in immune surveillance. However, cancer cells can evade immune detection by inhibiting macrophage phagocytosis through the interaction of their surface CD47 with SIRPα on macrophages. To counteract this immune evasion mechanism and promote phagocytosis, we developed miCD47 to downregulate CD47 expression on tumor cells (Fig. 2a). Similarly, three miRs were engineered to target murine CD47: mus-miCD47a, mus-miCD47b, and mus-miCD47c. Expression vectors pDMP-miCD47a/b/c were constructed for transfection into mouse CT26 and 3T3 cell lines. The results demonstrate that all miCD47 constructs significantly reduce CD47 expression in CT26 cells, but not in 3T3 cells (Fig. 2b), highlighting the efficacy and cancer cell selectivity of the designed miCD47s. Among them, mus-miCD47b exhibited the highest knockdown efficiency and was subsequently used to construct the vector pAAV-miCD47. In a subsequent co-culture experiment, transfection of CT26 cells with this vector significantly enhanced the cytotoxic activity of mouse macrophage RAW264.7 cells against CT26 cells (Fig. 2c). In an experimental study involving mice with CT26 tumors, treatment with rAAV-miNT and rAAV-miCD47 was administered (Fig. 2d). The findings demonstrate that rAAV-miCD47 treatment significantly inhibits tumor growth compared to rAAV-miNT (Fig. 2e). The therapeutic efficacy of rAAV-miPDL1 is further corroborated by tumor imaging (Fig. 2f), tumor size measurements (Fig. 2g), tumor weight assessments (Fig. 2g), and histological analysis via H&E staining of tumor tissues (Fig. 2h). The therapeutic impact of rAAV-miCD47 is also evidenced by a significant reduction in both the size and weight of the spleens (Fig. 2f and 2g). Preliminary evaluations of the biosafety of this treatment suggest minimal impact on the body weight of the mice (Additional file 1: Fig. S2a). This biosafety is primarily attributed to the selective knockdown of CD47 in tumor tissues, without affecting other healthy tissues (Fig. 2i), despite the widespread distribution of rAAV across all tissues (Additional file 1: Fig. S2b). Further indicators of the treatment’s biosafety include negligible effects on blood biochemical markers (WBC, RBC, PLT, HGB) (Additional file 1: Fig. S2c), hepatotoxicity markers (ALT, AST, ALP) (Additional file 1: Fig. S2d), kidney injury markers (BUN, Cr, UA) (Additional file 1: Fig. S2d), and the structural integrity of healthy tissues (Additional file 1: Fig. S2e). The data demonstrate the antitumor efficacy of rAAV-miCD47, highlighting its high tumor selectivity and biosafety in vivo.

**Fig. 2.**
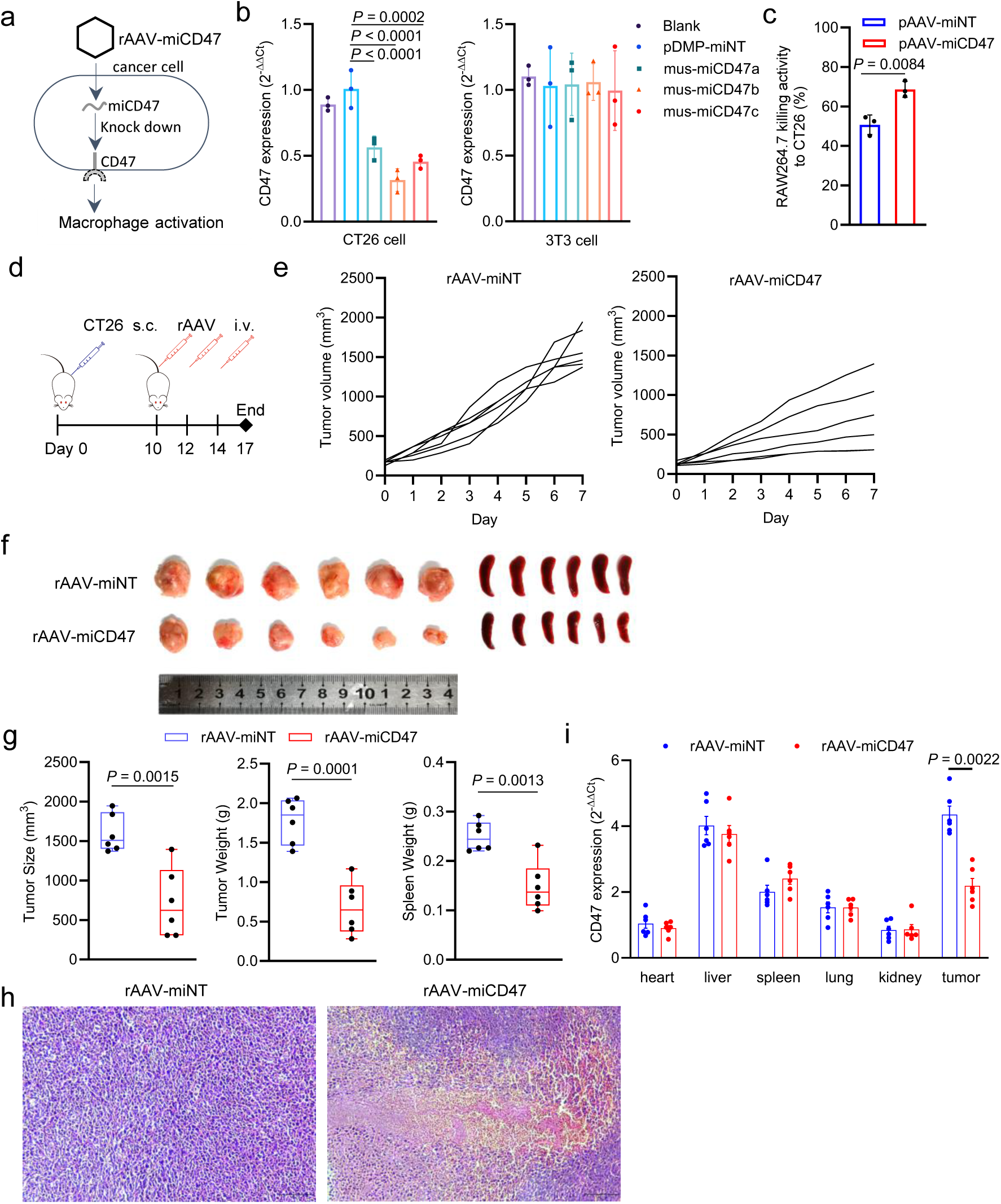
Treatment of CT26-xenografted mouse with rAAV-miCD47. **a** Schematics of inhibiting CD47 expression with rAAV-miCD47. **b** CD47 expression in CT26 and 3T3 cells with different transfection detected by qPCR (n=3 wells). **c** Killing activity of RAW264.7 cells against CT26 cells detected by CCK-8 reagent (n=3 wells). **d** Schematics of animal treatment (CT26 xenograft mice). s.c., subcutaneous injection; i.v., intravenous injection. **e** Tumor growth curve (n = 6 mice). **f** Tumor and spleen imaging. **g** Tumor volume, tumor weight, and spleen weight (n=6 mice). **h** Representative HE-staining tissue section of tumors. **i** Expression of CD47 in a variety of tissues detected by qPCR (n=6 mice).

### 3. PD-L1 and CD47 co-knockdown to treat mouse colon cancer

To enhance therapeutic outcomes through synergistic antitumor effects, we developed a new vector, pAAV-miPDL1-miCD47, capable of co-expressing miPDL1 and miCD47. This vector was packaged into an AAV and administered to mice. Mice bearing CT26 tumors received treatments of PBS, rAAV-miNT, rAAV-miPDL1, rAAV-miCD47, and rAAV-miPDL1-miCD47, respectively (Fig. 3a). The findings indicate that the rAAV-miPDL1-miCD47 treatment significantly inhibits tumor growth compared to treatments with rAAV-miPDL1 or rAAV-miCD47 alone (Fig. 3b). The synergistic antitumor effects of rAAV-miPDL1-miCD47 are further corroborated by tumor imaging (Fig. 3c), measurements of tumor size (Fig. 3d), tumor weight (Fig. 3d), and histological analysis via H&E staining of tumor tissues (Fig. 3e). The superior antitumor efficacy of rAAV-miPDL1-miCD47 is also evidenced by the significant reduction in spleen size and weight (Fig. 3c and 3d). Preliminary assessments of biosafety indicate that rAAV-miPDL1-miCD47 treatment has minimal impact on the body weight of mice (Additional file 1: Fig. S3a). The biosafety of the treatment is primarily attributed to the selective co-knockdown of PDL1 and CD47 in tumor tissues, while sparing healthy tissues (Fig. 3f), even though rAAV is distributed across all tissues (Additional file 1: Fig. S3b). The rAAV-miPDL1-miCD47 treatment demonstrates minimal impact on hematological parameters (WBC, RBC, PLT, HGB) (Additional file 1: Fig. S3c), hepatic function markers (ALT, AST, ALP) (Additional file 1: Fig. S3d), and renal function indicators (BUN, Cr, UA) (Additional file 1: Fig. S3d), as well as the structural integrity of healthy tissues (Additional file 1: Fig. S3e). These findings underscore the enhanced antitumor efficacy, high tumor specificity, and favorable biosafety profile of the rAAV-miPDL1-miCD47 treatment in vivo.

**Fig. 3.**
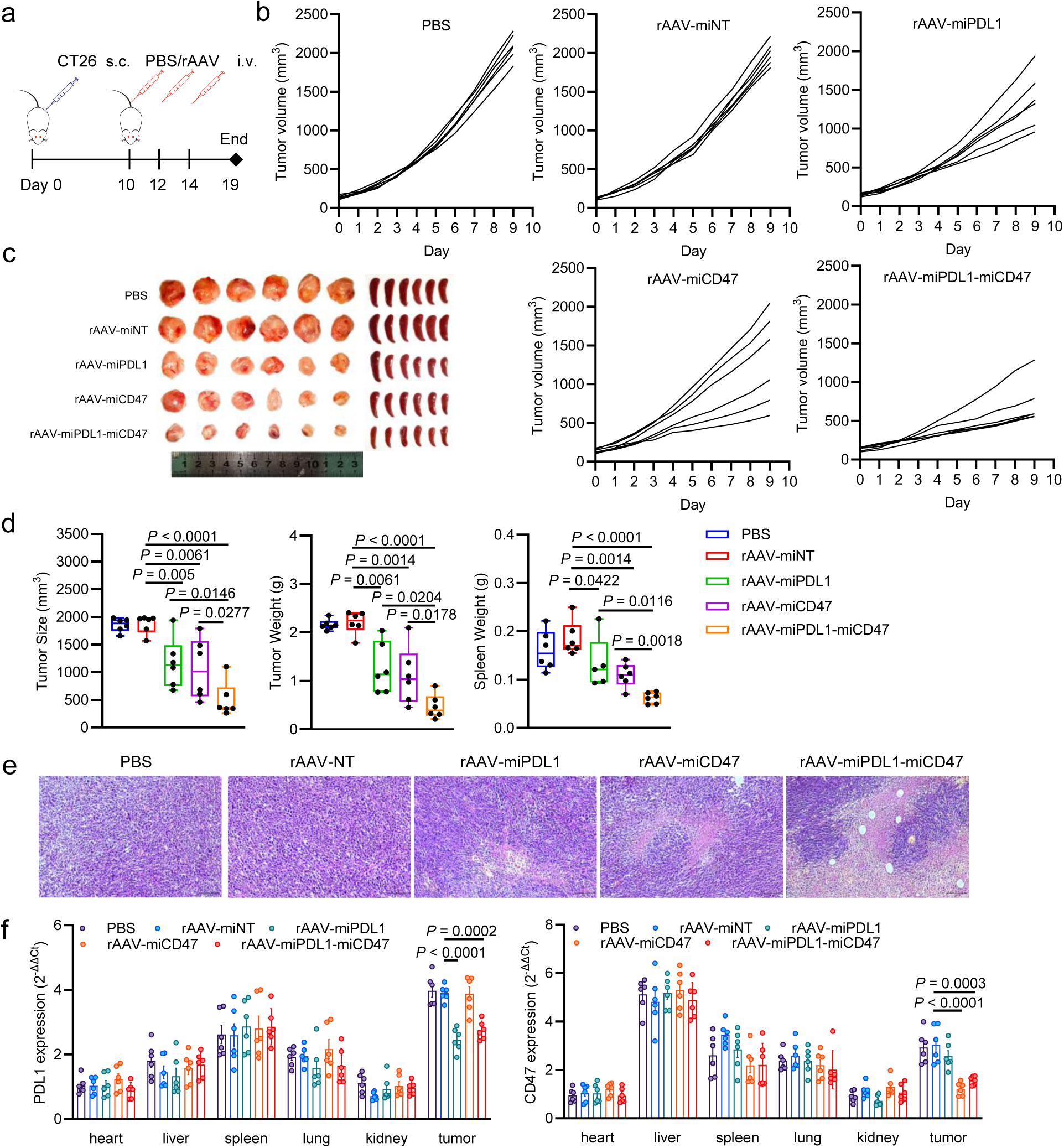
Treatment of CT26-xenografted mice using rAAV-miPDL1-miCD47. **a** Schematic representation of knocking down PD-L1 and CD47 expression with rAAV-miPDL1-miCD47. **b** PD-L1 and CD47 expression in CT26 and 3T3 cells following various transfections, as measured by qPCR (n=3 wells). **c** Killing activity of RAW264.7 cells against CT26 cells detected by CCK-8 reagent (n=3 wells). **d** Schematic overview of animal treatment protocol for CT26 xenograft mice, including subcutaneous (s.c.) and intravenous (i.v.) injections. **e** Tumor growth curve (n = 6 mice). **f** Imaging of the tumor and spleen. **g** Tumor volume, tumor weight, and spleen weight (n=6 mice). **h** Representative HE-staining tissue section of tumors. **i** Expression of CD47 levels in various tissues, as detected by qPCR (n=6 mice).

### 4. IL-15 over-expression to treat mouse colon cancer

The antitumor efficacy of immune cells, such as T cells, is frequently diminished due to chronic antigenic stimulation by tumors. Cytokines play a crucial role in enhancing the antitumor activity of immune cells, including T and NK cells. To further augment the antitumor efficacy of rAAV-miPDL1-miCD47, we investigated the expression of IL-15, a pivotal cytokine that activates both T and NK cells. Consequently, we constructed the vector pAAV-DMP-mIL15-WPRE (pAAV-mIL15) to facilitate the expression of IL-15 in cancer cells (Fig. 4a). This vector was subsequently employed to transfect mouse CT26 and 3T3 cells, and the IL-15 levels in the culture media were quantified using ELISA. The results indicate that IL-15 was successfully secreted by the pAAV-mIL15-transfected CT26 cells, but not by the pAAV-mIL15-transfected 3T3 cells (Fig. 4b), demonstrating the cancer cell-specific secretion of IL-15 by the vector. To assess the bioactivity of the secreted IL-15, the culture media from both cell types were mixed in a 1:1 ratio with CTLL2-specific medium to culture mouse T cell line CTLL2. The viability of CTLL2 cells was subsequently evaluated using the CCK-8 assay. The findings demonstrate that CTLL2 cells were effectively activated by the culture medium derived from pAAV-mIL15-transfected CT26 cells (Fig. 4c), thereby confirming the bioactivity of IL-15 secreted by the transfected cancer cells. Consequently, the vector was encapsulated into an rAAV-mIL15 for therapeutic application in mice (Fig. 4d). Mice bearing CT26 tumors received treatment with either rAAV-MCS or rAAV-mIL15 (Fig. 4d). The data reveal that rAAV-mIL15 treatment significantly inhibited tumor growth compared to rAAV-MCS (Fig. 4e). The therapeutic efficacy of rAAV-mIL15 is further corroborated by tumor imaging (Fig. 4f), as well as measurements of tumor size and weight (Fig. 4g), and histological analysis via H&E staining of tumor tissues (Fig. 4h). Additionally, the therapeutic effect of rAAV-mIL15 is evidenced by a marked reduction in the size and weight of the spleens (Fig. 4f and 4g). Preliminary assessments of biosafety suggest minimal impact on the body weight of treated mice (Additional file 1: Fig. S4a), which is primarily attributed to the selective overexpression of IL-15 in tumor tissues, with no significant expression in other healthy tissues (Fig. 4i), despite the widespread distribution of rAAV across all tissues (Additional file 1: Fig. S4b). The biosafety of this treatment is further demonstrated by its minimal impact on blood biochemical markers, including WBC, RBC, PLT, and HGB (Additional file 1: Fig. S4c). Additionally, the treatment exhibits negligible effects on hepatotoxicity markers, such as ALT, AST, and ALP Additional file 1: Fig. S4d), as well as on renal injury markers, including BUN, Cr, and UA (Additional file 1: Fig. S4d). Furthermore, the structures of healthy tissues remain unaffected (Additional file 1: Fig. S4e). Collectively, these data underscore the antitumor efficacy of rAAV-mIL15, highlighting its pronounced tumor selectivity and biosafety in vivo.

**Fig. 4.**
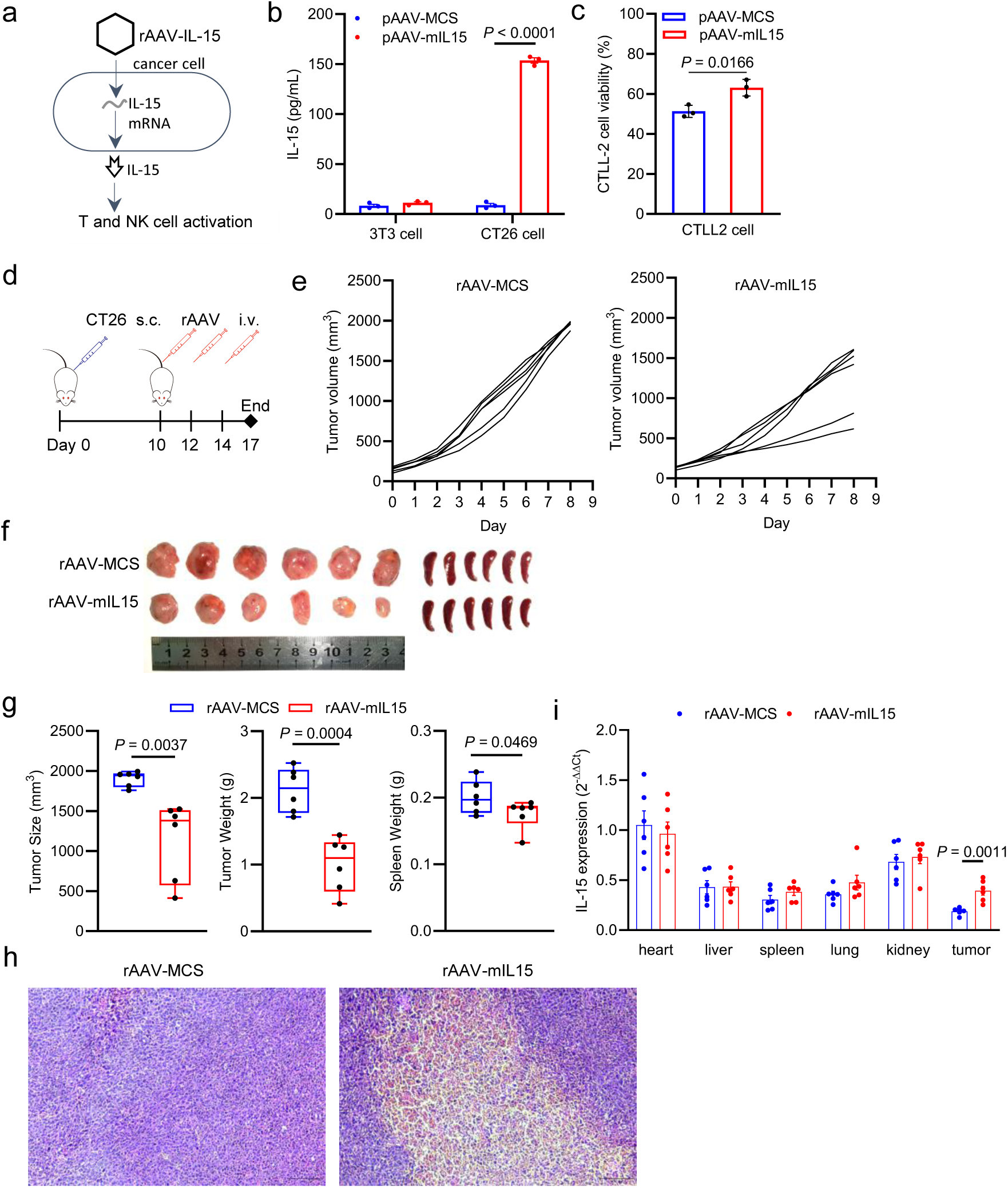
Treatment of CT26-xenografted mice using rAAV-mIL15. **a** Schematic representation f over expression of IL-15 with rAAV-mIL15. **b** IL-15 expression in CT26 cells and 3T3 cells detected by ELISA (n = 3 wells). **c** The CTLL2 cell viability in co-culture system detected by CCK8 reagent (n = 3 wells). **d** Schematic overview of animal treatment protocol for CT26 xenograft mice, including subcutaneous (s.c.) and intravenous (i.v.) injections. **f** Imaging of the tumor and spleen. **g** Tumor volume, tumor weight, and spleen weight (n=6 mice). **h** Representative HE-staining tissue section of tumors. **i** Expression of IL-15 levels in various tissues, as detected by qPCR (n=6 mice).

### 5. PD-L1 and CD47 co-knockdown plus IL-15 overexpression to treat mouse colon cancer

Upon confirming the antitumor efficacy of rAAV-mIL15, we proceeded to incorporate it into the rAAV-miPDL1-miCD47 construct by developing a new vector, pAAV-miPDL1-miCD47-mIL15, capable of co-expressing miPDL1, miCD47, and mIL15. This vector was subsequently packaged into AAV for administration in a murine model. Mice bearing CT26 tumors were treated with rAAV-MCS-NT, rAAV-miPDL1-miCD47, and rAAV-miPDL1-miCD47-mIL15, respectively (Fig. 5a). The findings demonstrate that the rAAV-miPDL1-miCD47-mIL15 treatment significantly inhibited tumor growth compared to rAAV-miPDL1-miCD47 alone (Fig. 5b). The synergistic antitumor effects of rAAV-miPDL1-miCD47-mIL15 were further corroborated by tumor imaging (Fig. 5c), tumor size (Fig. 5d), tumor weight (Fig. 5d), and histological analysis via H&E staining of tumor tissues (Fig. 5e). Additionally, the superior antitumor efficacy of rAAV-miPDL1-miCD47-mIL15 was evidenced by the marked reduction in spleen size and weight (Figs. 5c and 5d). Ultimately, the enhanced antitumor effects of rAAV-miPDL1-miCD47-mIL15 were reflected in the improved survival rates observed in treated mice compared to those receiving rAAV-miPDL1-miCD47 (Fig. 5f). Preliminary assessments of the biosafety profile of rAAV-miPDL1-miCD47-mIL15 indicated minimal impact on the body weight of the mice (Additional file 1: Fig. S5a). The biosafety of this approach is primarily attributed to the selective co-knockdown of PDL1 and CD47, along with the overexpression of IL-15 within tumor tissues, while sparing healthy tissues (Fig. 5g), despite the widespread distribution of rAAV across all tissues (Additional file 1: Fig. S5b). The rAAV-miPDL1-miCD47-mIL15 treatment demonstrates minimal impact on hematological parameters (WBC, RBC, PLT, HGB) (Additional file 1: Fig. S5c), hepatotoxicity markers (ALT, AST, ALP) (Additional file 1: Fig. S5d), and renal injury markers (BUN, Cr, UA) (Additional file 1: Fig. S5d), as well as on the structural integrity of healthy tissues (Additional file 1: Fig. S5e). Furthermore, the biosafety is supported by the absence of a systemic increase in serum IL-15 levels (Fig. 5h). These findings underscore the enhanced antitumor efficacy of rAAV-miPDL1-miCD47-mIL15, along with its high tumor selectivity and biosafety in vivo. The superior antitumor effect of rAAV-miPDL1-miCD47-IL15 was corroborated by an independent animal study (Additional file 1: Fig. S6).

**Fig. 5.**
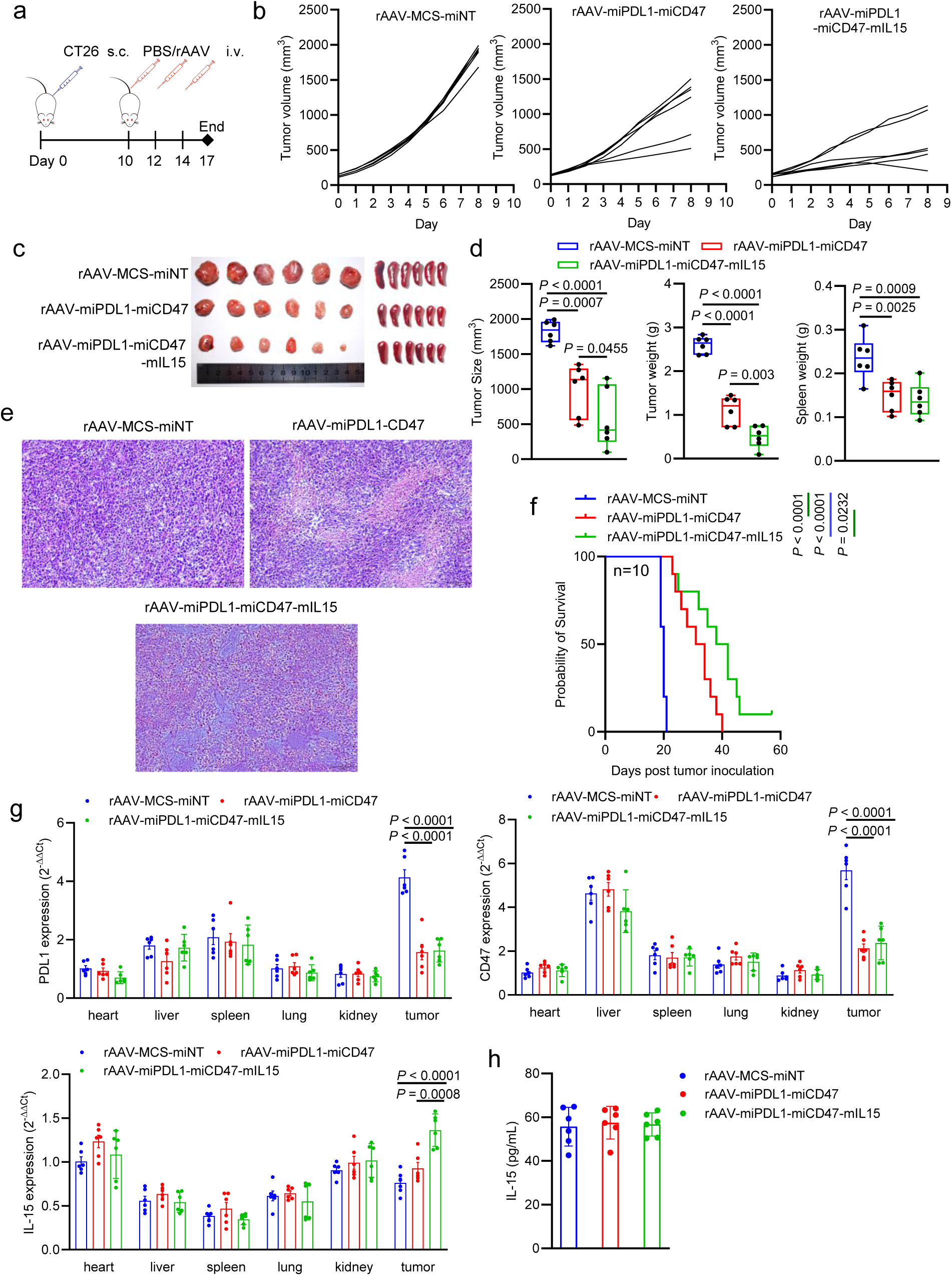
Treatment of CT26-xenografted mice using rAAV-miPDL1-miCD47-mIL15. **a** Schematic overview of animal treatment protocol for CT26 xenograft mice, including subcutaneous (s.c.) and intravenous (i.v.) injections. **b** Tumor growth curve (n = 6 mice). **c** Imaging of the tumor and spleen. **d** Tumor volume, tumor weight, and spleen weight (n=6 mice). **e** Representative HE-staining tissue section of tumors. **f** Kaplan-Meier survival curve (n = 10 mice) **g** Expression of PDL1, CD47, and IL-15 levels in various tissues, as detected by qPCR (n=6 mice). **h** IL-15 level in serum, as detected by ELISA (n = 6 mice).

### 6. Similar immunogene therapy to treat human cancer cells

To further explore the potential clinical application of this strategy in human patients, we designed analogous miRs targeting human PD-L1 and CD47 genes. We constructed pAAV-miPDL1/miCD47 vectors to achieve knockdown of human PD-L1 and CD47 expression and utilized these vectors to transfect human lung cancer cell line A549, human liver cancer cell line HepG2, human colon cancer cell line HCT116, and human embryonic lung fibroblast cell line MRC5. The results demonstrated a significant reduction in PD-L1 expression in the cancer cell lines, A549 and HepG2, but not in the normal cell line MRC5 (Additional file 1: Fig. S7a). Among the miRs tested, homo-miPDL1a exhibited the highest knockdown efficiency and was subsequently employed to activate human T cells. The human colon cancer cell line HCT116 was transfected with pAAV-DMP-homo-miPDL1a (pAAV-miPDL1) and co-cultured with human T cell line Jurkat. Following co-culture, the suspended Jurkat cells were collected, and the expression levels of IL-2 and IFN-γ were assessed. The findings demonstrate that the expression of the two genes, indicative of T cell activation, was significantly upregulated (Additional file 1: Fig. S7b), suggesting that the Jurkat cells were activated following PD-L1 knockdown in HCT116 cells (Additional file 1: Fig. S7a). This observation is consistent with the marked decrease in HCT116 cell viability post-co-culture (Additional file 1: Fig. S7c). Subsequently, similar transfection procedures and human macrophage activation assays were conducted using miRs targeting the human CD47 gene. The results indicated that human CD47 expression was significantly reduced by all three miCD47 constructs in two human cancer cell lines (A549 and HCT116), but not in the human normal cell line MRC5 (Additional file 1: Fig. S8a). Among these, homo-miCD47c exhibited the highest knockdown efficiency and was therefore utilized for human macrophage activation. Human macrophages were prepared by stimulating THP-1 cells with PMA and LPS (Additional file 1: Fig. S8b). The prepared macrophages were subsequently utilized to treat pAAV-DMP-homo-miCD47c-transfected human A549 and HCT116 cells. The results demonstrated a significant cytotoxic effect on both cancer cell lines by the macrophages (Additional file 1: Fig. S8c), suggesting that macrophage activation was achieved through the knockdown of CD47 in the cancer cells. Subsequently, human MRC5 and HCT116 cells were transfected with pAAV-MCS/pAAV-hIL15, and IL-15 expression was assessed via qPCR. The findings indicated that IL-15 expression was exclusive to HCT116 cells (Additional file 1: Fig. S9a and S9b). The pAAV-MCS/pAAV-hIL15-transfected HCT116 cells were then co-cultured with PMA and PHA-activated Jurkat cells. The results revealed that the transfection with pAAV-hIL15 significantly enhanced the susceptibility of HCT116 cells to cytotoxicity by Jurkat cells (Additional file 1: Fig. S9c). These findings suggest that this strategy may be applicable for the treatment of human cancers.

## Discussion

In recent years, immunotherapy has emerged as a significant cancer treatment modality due to its high therapeutic efficacy, particularly therapies based on immune checkpoint inhibitors (ICIs), which can achieve sustained remission in responsive patients. However, these therapies are still confronted with severe toxic effects in clinical settings [28, 45–47], notably various immune-related adverse events (irAEs) that affect healthy tissues due to on-target, off-tumor side effects. Consequently, there is a pressing need for the development of safer immunotherapies targeting immune checkpoints (ICs). Since the mechanisms of ICs are utilized by both normal and tumor cells to evade immune cell attacks, the on-target, off-tumor toxicity associated with antibody-based ICIs appears to be an inherent challenge, indicating that overcoming this type of autoimmune toxicity using the current ICI strategies is impractical. In response to this issue, we explored an alternative approach. In this study, we developed a novel immunogene therapy that activates immune cells by selectively inhibiting the expression of known ICs in tumor cells. We demonstrate the in vivo antitumor efficacy and safety of this strategy.

PD-1 and its ligand, PD-L1, represent the most extensively utilized immune checkpoints in contemporary immunotherapy. Numerous antibody drugs targeting PD-1/PD-L1 have been widely implemented in clinical settings. Nevertheless, the current ICIs based on PD-1/PD-L1 inhibitors face challenges due to severe side effects derived from their mechanisms of action. Although PD-1 and PD-L1 function as a pair of co-inhibitory molecules [48], PD-1 is primarily expressed on activated T cells, whereas PD-L1 is expressed on both normal and tumor cells. Despite sharing a common therapeutic principle, PD-1/PD-L1 inhibitors exhibit varying therapeutic effects [49]. PD-1 inhibitors are predominantly IgG4 antibodies, which possess limited structural stability and require modifications to enhance stability and reduce adverse reactions [50]. In contrast, PD-L1 inhibitors are primarily IgG1 antibodies characterized by a prolonged half-life and robust stability. PD-1 inhibitors may increase the risk of autoimmune reactions by binding to both PD-L1 and PD-L2 on immune cells. Conversely, PD-L1 inhibitors can result in stronger overall immune efficacy by binding to PD-L1 on both cancer and antigen-presenting cells [51, 52]. Although PD-L1 antibodies exhibit a lower incidence of irAEs compared to PD-1 antibodies [52–55], they are still significantly challenged by on-target off-tumor effects, due to lack of tumor specificity. These effects can lead to severe autoimmune diseases in extreme cases [26, 56], and increase the risk of infection by affecting B cell function [57]. Furthermore, PD-L1 antibodies can be substantially neutralized by PD-L1-bearing exosomes derived from tumor cells [58–60]. In this study, we aimed to knock down PD-L1 expression on cancer cells. Our findings demonstrate that rAAV-miPDL1 exhibits significant antitumor effects and high safety by specifically targeting and reducing PD-L1 expression in tumors. This approach not only replicates the antitumor efficacy of PD-L1 inhibitors but also mitigates their off-tumor side effects. Moreover, this strategy can inhibit the release of PD-L1-bearing exosomes by tumor cells, which can circulate in the body and cause systemic immunosuppression by inhibiting T cells and DCs in the draining lymph nodes and spleen [61–65].

Innate immunity plays a crucial role in the surveillance and elimination of abnormal cells. For instance, the activation of macrophages, NK cells, and group 2 innate lymphoid cells (ILC2) has been extensively investigated for cancer treatment [66, 67]. As a pivotal immune checkpoint in innate immunity, the CD47-SIRPα interaction has been targeted to activate innate immune cells through the use of CD47 inhibitors, such as antibodies [68, 69]. Mechanistically, CD47 inhibitors promote macrophage phagocytosis by disrupting the interaction between CD47 on cancer cells and SIRPα on macrophages [70]. CD47 antibodies can effectively reprogram pro-tumor M2 macrophages into anti-tumor M1 macrophages [71–73]. Additionally, inhibiting the CD47-SIRPα interaction also stimulates NK cells to target cancer cells [74, 75]. However, drug development in this domain faces challenges due to the low maximum tolerated dose and limited therapeutic efficacy, primarily because of significant side effects [76]. These side effects largely arise from the toxicity of CD47 inhibitors to healthy cells, as CD47 is ubiquitously expressed as a “don”t-eat-me” signal.. The therapeutic potential of CD47 monoclonal antibodies is significantly constrained by their pronounced toxicity to erythrocytes, which can result in anemia [77]. Consequently, similar on-target off-tumor effects continue to impede the clinical application of CD47 inhibitors. There is an urgent need for strategies that maximize the pro-phagocytosis effects of anti-CD47 antibodies while minimizing adverse reactions. In this study, we aimed to specifically knock down CD47 expression in tumor cells. Our findings demonstrate that rAAV-miCD47 exhibits significant antitumor effects and high safety by specifically targeting CD47 expression in tumors. This immunogene therapy holds promise for facilitating the clinical application of CD47 inhibition in the near future.

Immune stimulators play a crucial role in effective cancer therapy, and cytokines and chemokines with immune-stimulating properties have been extensively investigated in cancer treatment [78]. For instance, IL-15 has garnered increasing interest in this field due to its unique function. IL-15 was identified in 1994 [79, 80] and shares numerous biological activities with IL-2, such as the stimulation of T cell proliferation, the induction of cytotoxic lymphocyte and memory phenotype CD8+ T cell generation, the promotion and maintenance of natural killer (NK) cell proliferation, and the induction of B cell immunoglobulin synthesis [36, 81]. Despite these similarities, IL-2 and IL-15 exhibit distinct in vivo functions, particularly in the context of adaptive immune responses. Both cytokines transmit signals through shared receptor subunits; however, IL-15 utilizes a unique trans-presentation mechanism via IL-15Rα, which ensures its specificity and function. This distinction renders IL-15 crucial for long-term immune response and cell survival, whereas IL-2 predominantly facilitates the activation and regulation of immune responses [82, 83]. Notably, IL-2 is essential for the development and maintenance of regulatory T cells (Treg), which are intimately linked to activation-induced cell death (AICD) [84]. Conversely, IL-15 plays a pivotal role in sustaining the persistence of NK and memory CD8+ T cells. Unlike IL-2, IL-15 does not mediate AICD; instead, it inhibits IL-2-induced AICD [85]. IL-15 emerges as a promising cytokine for cancer immunotherapy; however, its clinical application is constrained by dosage limitations and off-target toxicity beyond the tumor site [86]. In this study, we endeavored to enhance the expression of IL-15 specifically within tumor tissues utilizing recombinant adeno-associated virus encoding murine IL-15 (rAAV-mIL15). The findings demonstrate that rAAV-mIL15 exerts significant antitumor effects while maintaining a high safety profile, attributable to the localized expression of IL-15 within the tumor microenvironment. The safety of rAAV-mIL15 is further corroborated by the absence of alterations in serum IL-15 levels, thereby mitigating the risk of systemic toxicity.

The immune system’s ability to eradicate tumors is multifaceted, and various immunotherapies possess distinct advantages and limitations. Combination therapies are increasingly regarded as promising approaches to enhance therapeutic efficacy. Notably, the antitumor activity of immune checkpoint inhibitors (ICIs) when used as monotherapies has been insufficient in significantly improving overall survival [87]. Consequently, the clinical adoption of combination therapies involving different ICIs has been pursued to enhance therapeutic outcomes [87–89]. For example, the combination of CTLA-4 and PD-1/PD-L1 antibodies are used to improve the therapeutic efficacy by integrating two ICIs with different mechanisms [90, 91]. However, this approach merely represents a combination of two separate exogenous antibodies [92], rather than a single drug targeting multiple pathways. In contrast, immunogene therapy offers a unified strategy with a single agent targeting multiple pathways. This approach effectively integrates the antitumor effects of various ICIs targeting different immune checkpoints, such as PD-L1 and CD47 [88]. By simultaneously inhibiting multiple immune checkpoints on a single cancer cell, this strategy activates diverse immune responses, thereby maximizing the potential for complete tumor eradication. However, the combination of ICI antibodies alone may have a reduced likelihood of achieving such comprehensive outcomes. In this study, the rAAV vectors rAAV-miPDL1 and rAAV-miCD47, which target adaptive and innate immunity respectively, were combined into a single vector, rAAV-miPDL1-miCD47. This vector is capable of co-expressing miRs that target both PD-L1 and CD47 on cancer cells. The findings demonstrated that rAAV-miPDL1-miCD47 exhibited superior therapeutic efficacy compared to either rAAV-miPDL1 or rAAV-miCD47 alone. Furthermore, it is feasible to incorporate additional cassettes expressing miRs targeting multiple ICs into a single AAV particle to achieve enhanced synergistic therapeutic effects in future applications. Beyond IC-targeting miRs, cassettes encoding cytokines with strong immunostimulatory functions, such as IL-15 used in this study, can also be integrated into a single AAV particle, provided the total inserted DNA remains within the AAV packaging capacity (up to 4.7 kilobases). Consequently, the ability to integrate diverse immunotherapeutic mechanisms into a single AAV vector represents a significant advantage of this immunogene therapy approach.

The adverse effects of current ICIs on healthy tissues primarily arise from on-target, off-tumor side effects. Consequently, achieving selective targeting of tumor cells in vivo is crucial for the safety of this immunogene therapy. Establishing tumor cell specificity is therefore essential. To address this, we systematically assessed the expression levels of PD-L1, CD47, and IL-15 in cultured cells and various tissues using qPCR, which offers high detection sensitivity. Our qPCR analysis of multiple tissues, including the heart, liver, spleen, lung, kidney, and tumors from each mouse across all animal treatment experiments, revealed that the expression of the two target ICs, PD-L1 and CD47, was significantly reduced only in tumor tissues. Their expression levels in all examined healthy tissues remained unaffected by AAV treatment. Similarly, IL-15 was overexpressed exclusively in tumor tissues, with no alteration in expression observed in healthy tissues. These findings underscore the high tumor selectivity of this immunogene therapy, effectively mitigating the off-tumor effects associated with current antibody-based ICIs. The tumor selectivity of this immunogene therapy is consistent with its favorable biosafety profile, as evidenced by the absence of significant effects on all assessed safety parameters, including body weight, blood cell composition, liver and kidney toxicities, and histopathological evaluations.

In this study, we employed adeno-associated virus (AAV) to deliver the therapeutic DNA into cells in vivo, thereby leveraging the advantages of AAV. AAV is characterized by a high transfection capacity across all in vivo cells, encompassing both non-dividing cells (e.g., normal cells) and dividing cells (e.g., cancer cells). AAV is recognized for its superior safety profile in current human gene therapy, with several AAV-based drugs already approved for the treatment of human genetic diseases in clinical settings [93]. Furthermore, AAV can now be commercially produced on a large scale by specialized contract manufacturing organizations (CMOs). In our study, we utilized a dose 100-fold lower (10^12^/kg body weight) than that typically used for treating human genetic diseases (10^14-15^/kg body weight) [93], thereby enhancing the safety and affordability of this immunogene therapy for future clinical applications. This study demonstrated that AAV is capable of distributing across all tissues with varying levels of abundance, exhibiting a relatively higher distribution within tumors. This finding aligns with extensive characterizations in our previous studies [42–44]. AAV itself is characterized by high safety, as evidenced by transfection with rAAVs used as negative controls, including rAAV-MCS, rAAV-miNT, and rAAV-MCS-miNT. These rAAVs were administered to mice at equivalent doses and did not produce significant effects on several key safety indices, such as body weight, blood cell counts, and liver and kidney toxicities. Based on this foundation, all therapeutic rAAVs, including rAAV-miPDL1, rAAV-miCD47, rAAV-mIL15, rAAV-miPDL1-miCD47, and rAAV-miPDL1-miCD47-mIL15, similarly exhibited no significant impact on these key safety indices. The safety profile was further corroborated by histological examination of various tissues from all mice using HE staining of tissue sections, which revealed no significant pathological effects on healthy tissues following administration of any rAAVs.

AAVs exhibit a broad natural infection profile in humans [94]. Consequently, there is a prevalent pre-existing humoral immunity to AAVs in the human population [94, 95]. Currently, AAV-based immunogene therapy faces challenges due to pre-existing anti-AAV antibodies and the need for repeated drug administration. However, several FDA-approved clinical agents have demonstrated significant efficacy in reducing or even eliminating anti-AAV antibodies, including neutralizing antibodies (NAbs) [96]. These agents include the neonatal Fc receptor (FcRn) antibody [97], IgG-cleaving endopeptidase or IgG-degrading enzyme (such as IdeS [98]; IdeZ [99], IceMG [100]), other IgG degrader (such as the dual-specific IgG degrader BHV-1300 of Biohaven)[101], and CD20 antibody [102, 103]. A recent study conducted by Wilson’s lab demonstrated that gene delivery to the liver and heart via systemic AAV gene therapy is feasible in mice and nonhuman primates by reducing pre-existing NAb levels using FcRn-inhibiting monoclonal antibodies [97]. Several clinical trials are currently being conducted to assess the clearance of pre-existing antibodies against AAVs, such as rAAVrh74 and AAV8, using IdeS. This approach aims to facilitate AAV-based gene therapies for genetic disorders, including Duchenne muscular dystrophy (NCT06241950) and Crigler-Najjar syndrome (NCT06518005). Consequently, the administration and repeated administration of these immunogene therapy-related AAVs to patient with various cancers is anticipated to become feasible in the near future.

## Conclusion

In this study, we have developed an innovative immunogene therapy to cancer, achieved through construction of therapeutic gene expression cassettes that are specifically activated by the intracellular NF-κB activity. These cassettes are designed to selectively express miRs targeting ICs such as PD-L1 and CD47, as well as the cytokine IL-15, within cancer cells. We have demonstrated their therapeutic efficacy and safety in murine models of colon cancer by systematically delivering them to tumors using AAV vectors. Consequently, we propose a novel immunotherapy strategy that offers an alternative to current antibody-based ICIs. This strategy is advantageous in mitigating the toxicities associated with antibody-based ICIs, which often arise from on-target, off-tumor side effects. More importantly, this approach effectively integrates the antitumor effects of various ICIs targeting different ICs, such as PD-L1 and CD47. Furthermore, it enhances the synergetic therapeutic efficacy of inhibiting multiple by facilitating tumor-specific overexpression of immunostimulatory cytokines like IL-15. These benefits of immunogene therapy are unattainable with current antibody-based immunotherapies. Therefore, this immunogene therapy offers a promising avenue for cancer treatment.

## Supplementary Information

The online version contains supplementary material available online. Additional file 1.

## Funding

This investigation was mainly funded by the project of the National Natural Science Foundation of China (62371126), and partially supported by the National Key Research and Development Program of China (2022YFA1205802) and the Suzhou Municipal Science and Technology Plan Funding Project (SY202201).

## Data Availability

All data generated or analyzed during this study are included in this published article.

## Declarations

### Ethics approval and consent to participate

All animal experiments were performed under protocols approved by the Animal Experimentation and Ethics Committee of Southeast University.

### Consent for publication

All authors of this study agreed to publish.

### Competing interests

The authors declare that they have no competing interests.

## Supplementary information

**Table S1.**
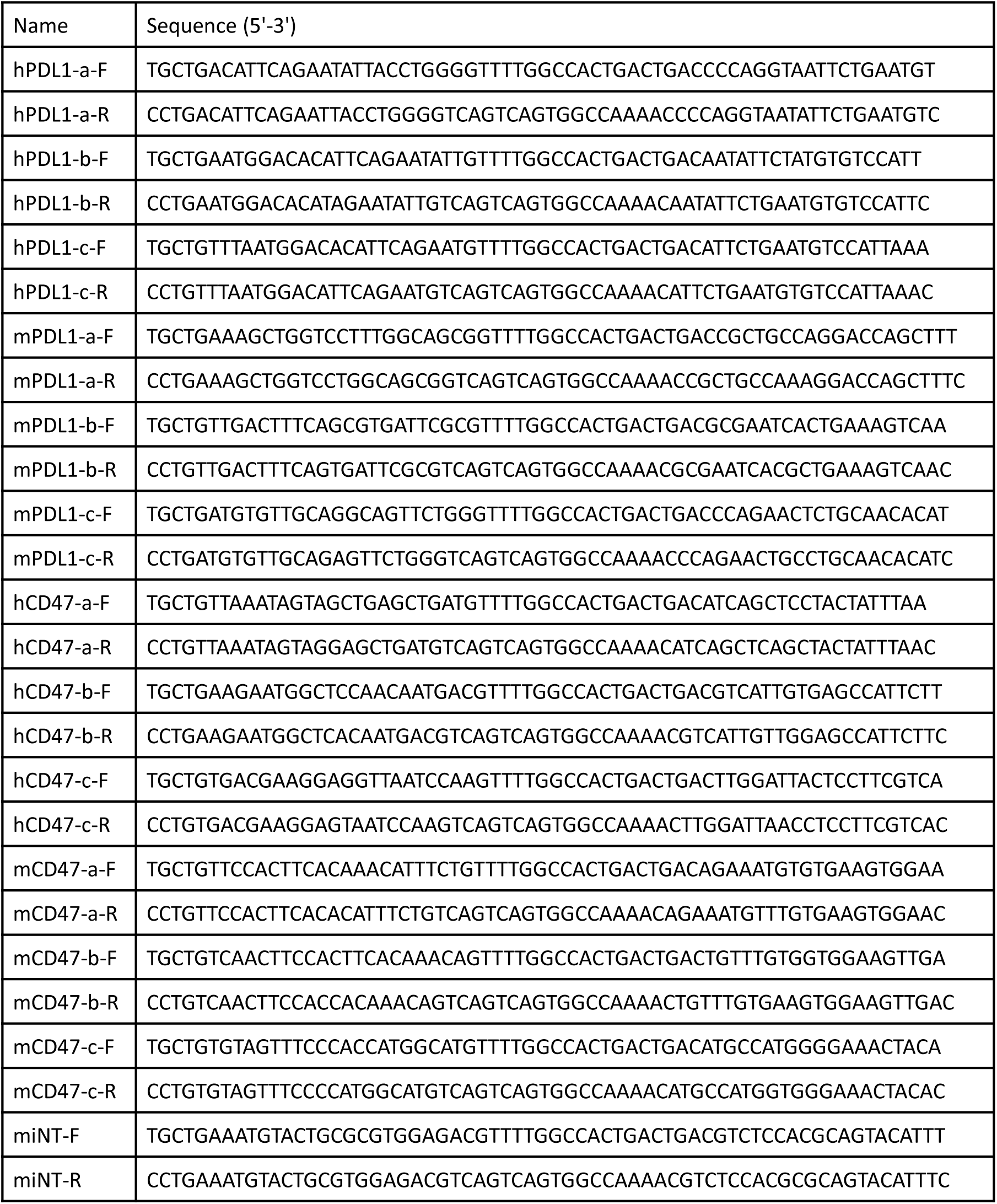
Oligonucleotides sequences used to construct miRNA expression vector.

**Table S2.**
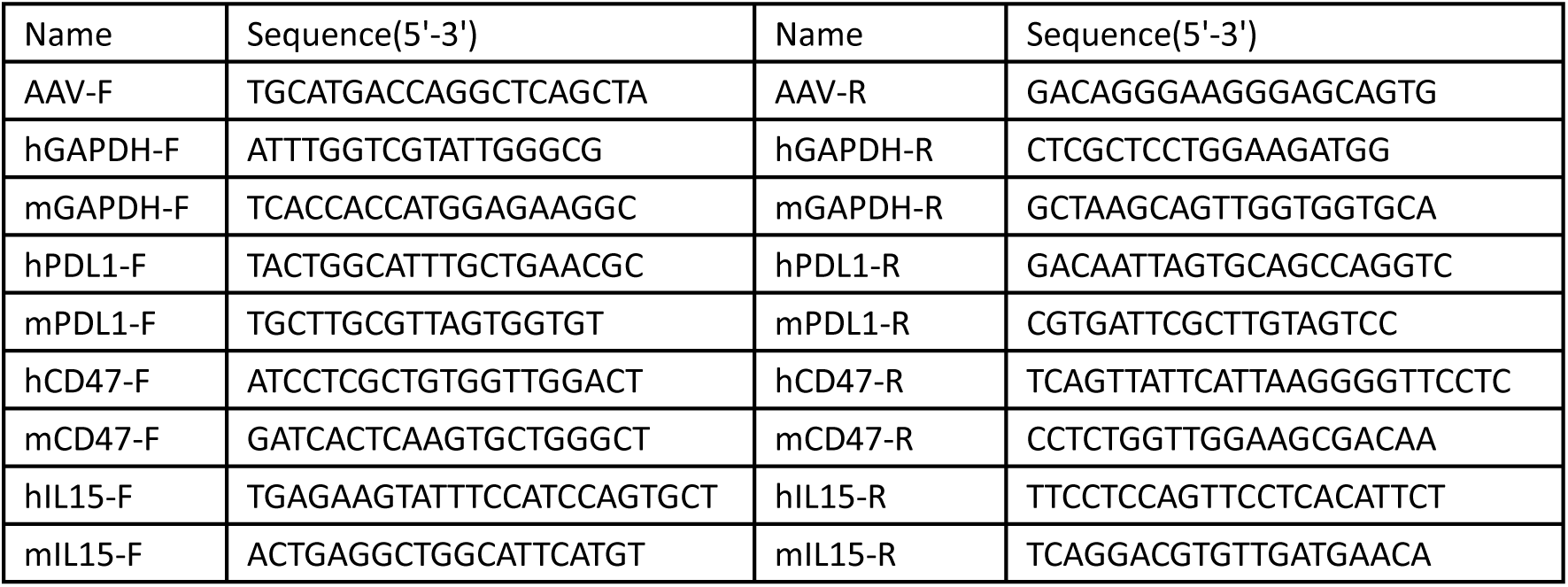
Primer sequences used for qPCR.

**Fig. S1.**
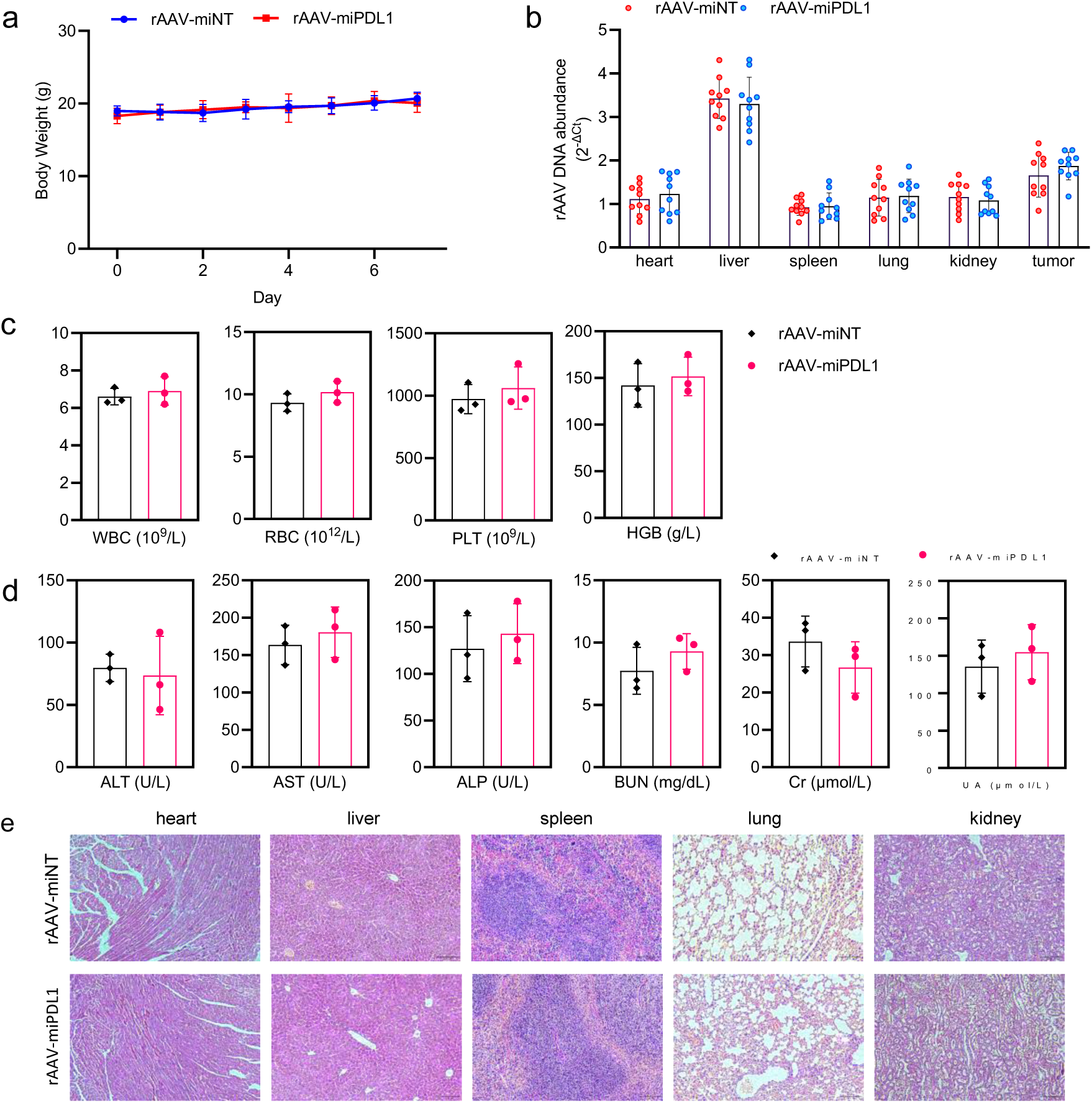
Treatment of CT26-xenografted mouse with rAAV-miPDL1. **a** Average body weight (n=10 mice). **b** Distribution of viral DNA in tissues. **c** Routine blood test (n=3 mice). **d** Serum biochemical indices detection (n=3 mice). **e** Representative images of H&E-stained tissue sections of a variety of tissues (heart, liver, spleen, lung, and kidney).

**Fig. S2.**
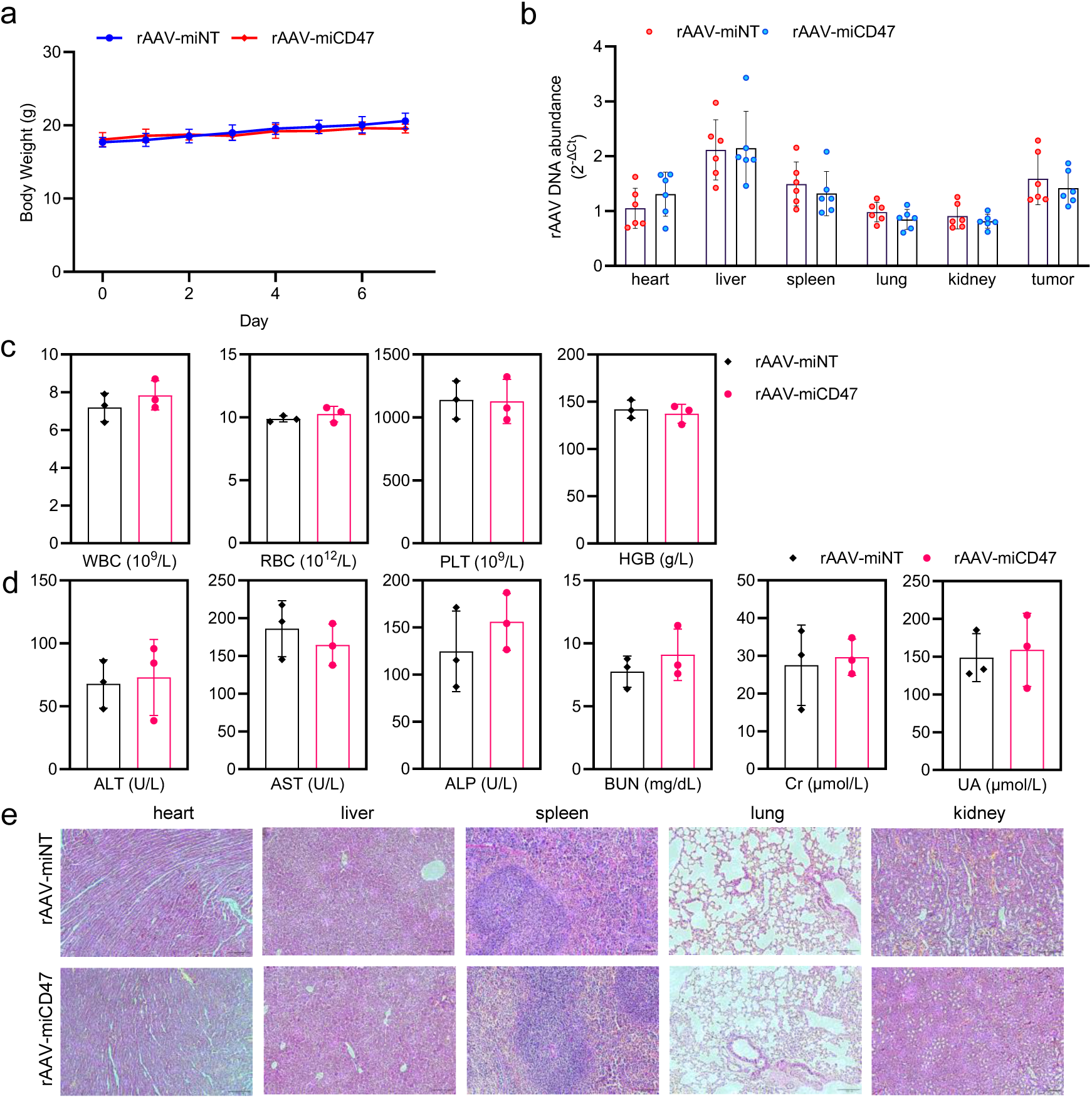
Treatment of CT26-xenografted mouse with rAAV-miCD47. **a** Average body weight (n=6 mice). **b** Abundance of AAV genomic DNA in a variety of tissues detected by qPCR (n=6 mice). **c** Routine blood test (n=3 mice). **d** Serum biochemical indices detection (n=3 mice). **e** Representative images of H&E-stained tissue sections of a variety of tissues (heart, liver, spleen, lung, and kidney).

**Fig. S3.**
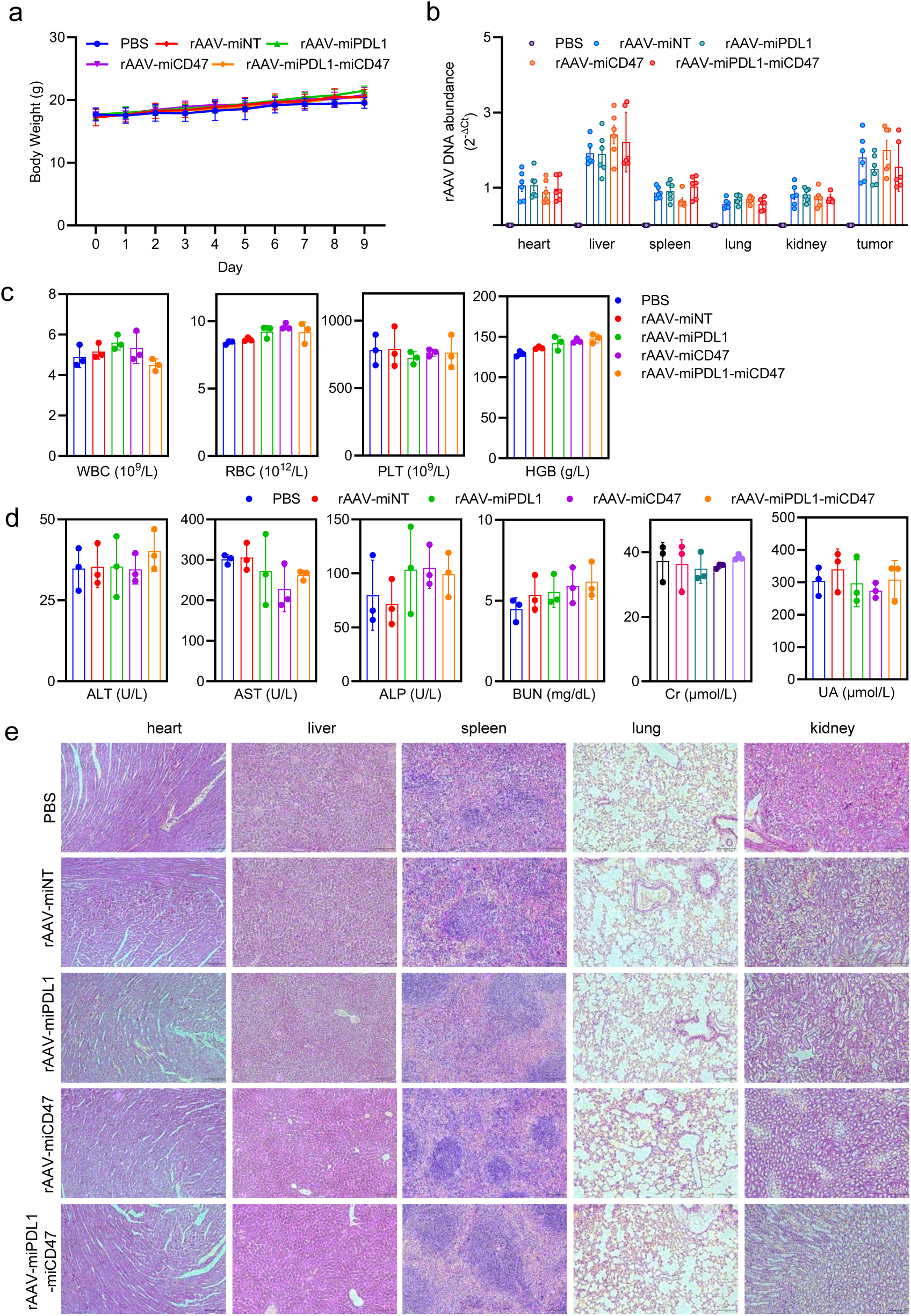
Treatment of CT26-xenografted mouse with rAAV-miPDL1-miCD47. **a** Average body weight. **b** Abundance of AAV genomic DNA in a variety of tissues detected by qPCR (n=6 mice). **b** Expression of PDL1 mRNA in tissues. **c** Routine blood test (n=3 mice). **d** Serum biochemical indices detection (n=3 mice). **e** Representative images of H&E-stained tissue sections of a variety of tissues (heart, liver, spleen, lung, and kidney).

**Fig. S4.**
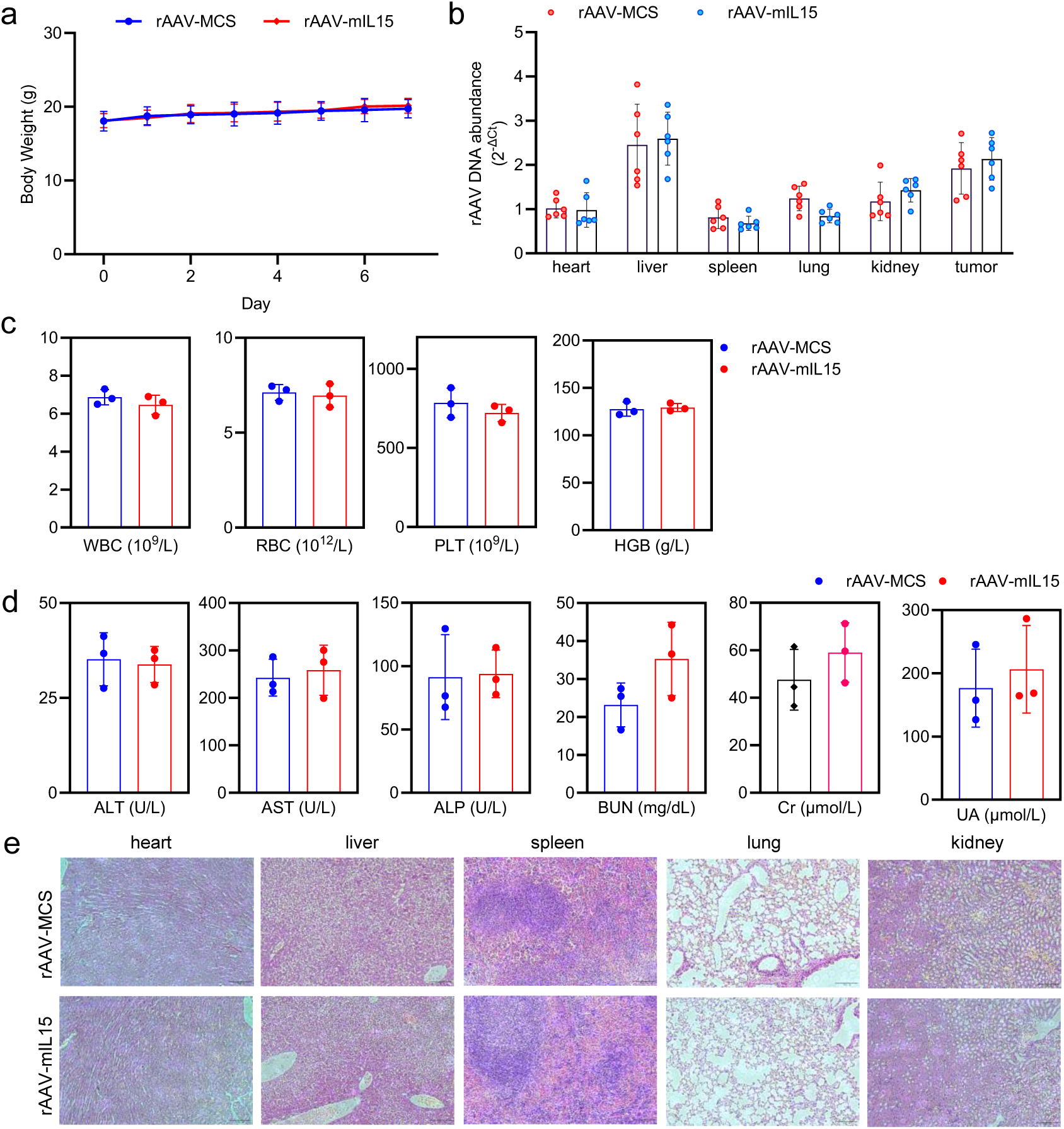
Treatment of CT26-xenografted mouse with rAAV-mIL15. **a** Average body weight. Data are presented as mean ± SD, n = 6 mice. **b** Abundance of AAV genomic DNA in a variety of tissues detected by qPCR (n=6 mice). **c** Routine blood test (n=3 mice). **d** Serum biochemical indices detection (n=3 mice). **e** Representative images of H&E-stained tissue sections of a variety of tissues (heart, liver, spleen, lung, and kidney).

**Fig. S5.**
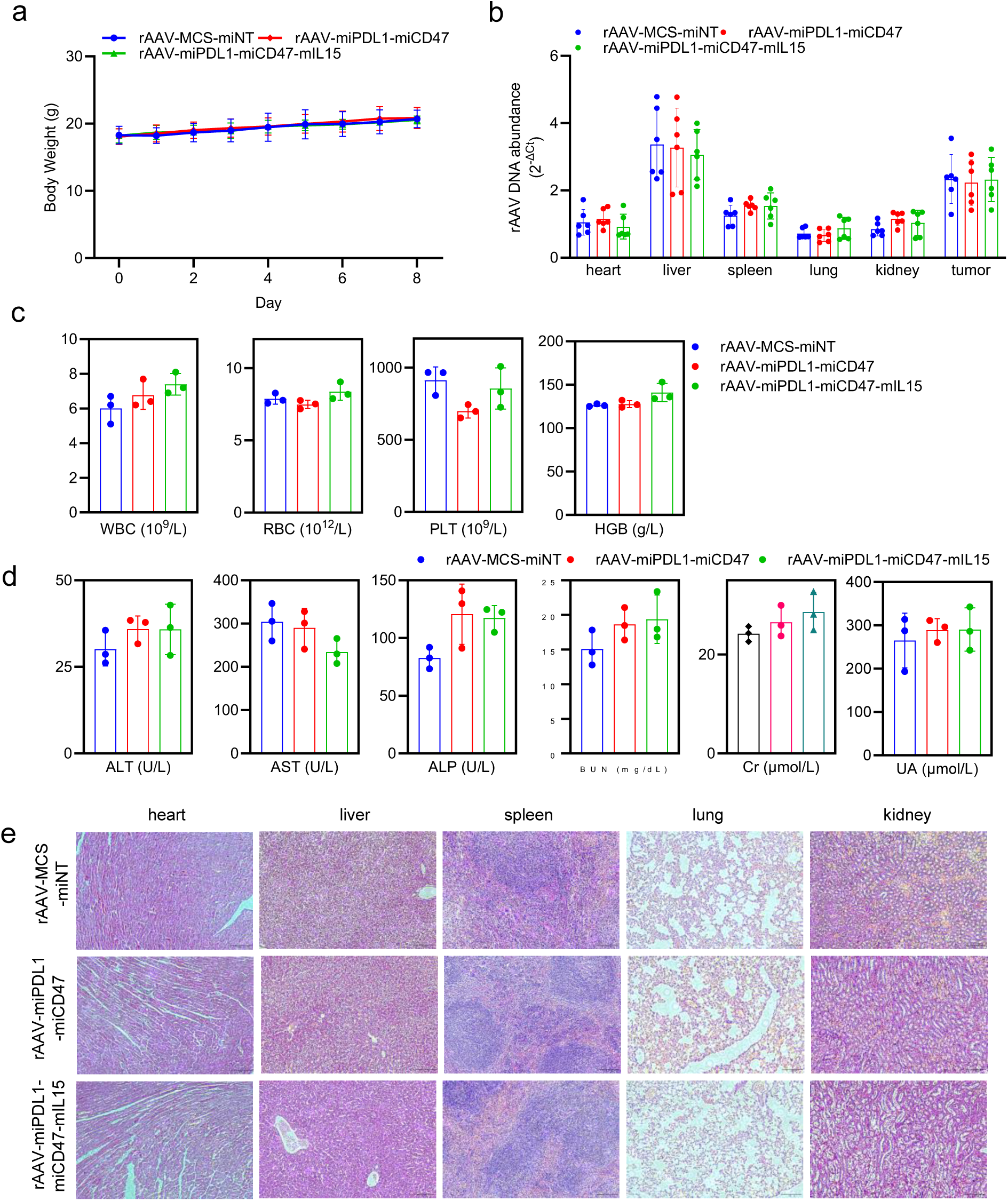
Treatment of CT26-xenografted mouse with rAAV-miPDL1-miCD47-mIL15. **a** Average body weight (n=3 mice). **b** Abundance of AAV genomic DNA in a variety of tissues detected by qPCR (n=6 mice). **c** Routine blood test (n=3 mice). **d** Serum biochemical indices detection (n=3 mice). **e** Representative images of H&E-stained tissue sections of a variety of tissues (heart, liver, spleen, lung, and kidney).

**Fig. S6.**
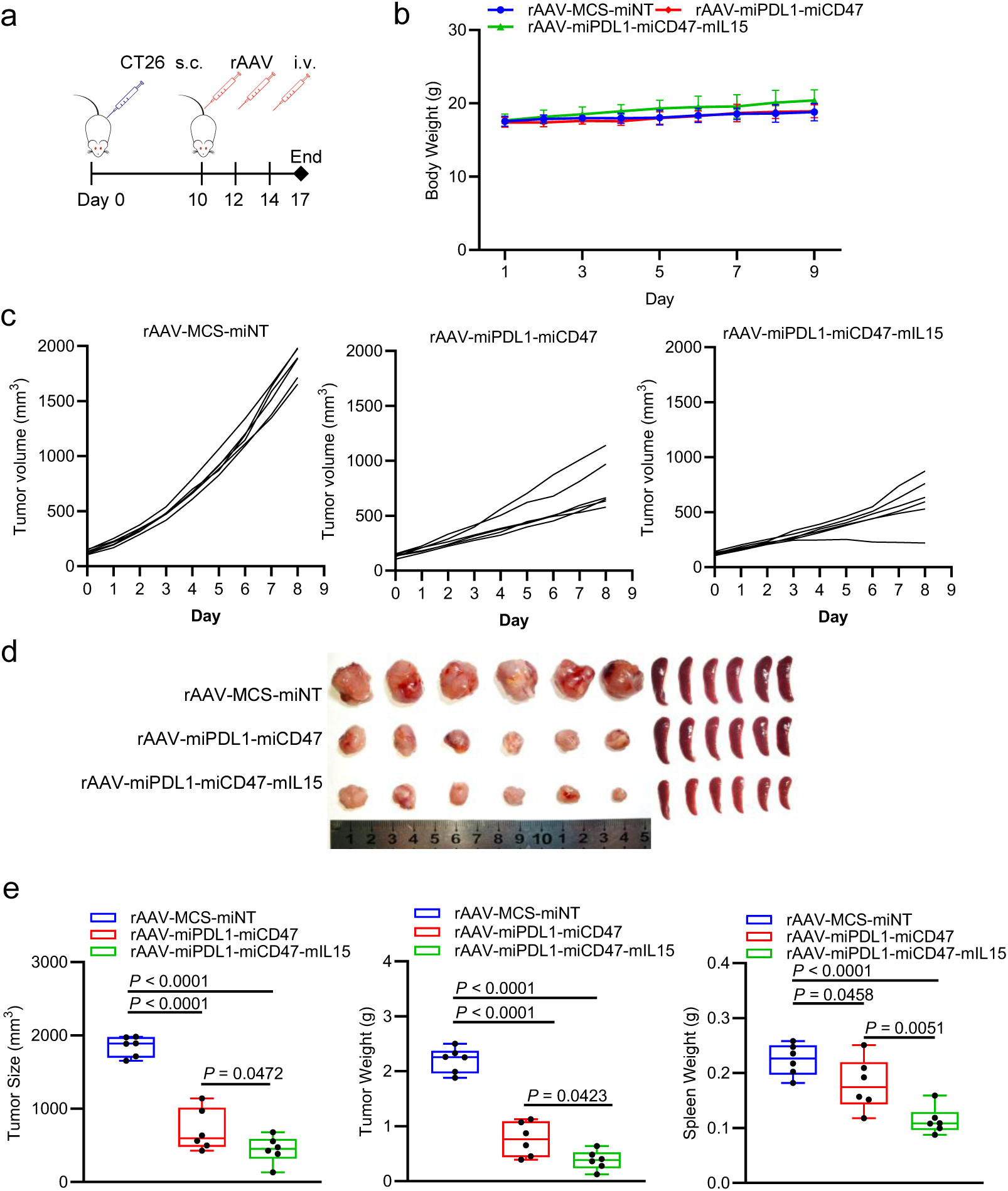
Treatment of CT26-xenografted mouse with rAAV-miPDL1-miCD47-mIL15. **a** Schematics of animal treatment (CT26 xenograft mice). s.c., subcutaneous injection; i.v., intravenous injection. **b** Average body weight (n=6 mice). **c** Tumor growth curve (n = 6 mice). **d** Tumor and spleen imaging. **e** Tumor volume, tumor weight, and spleen weight (n = 6 mice).

**Fig. S7.**
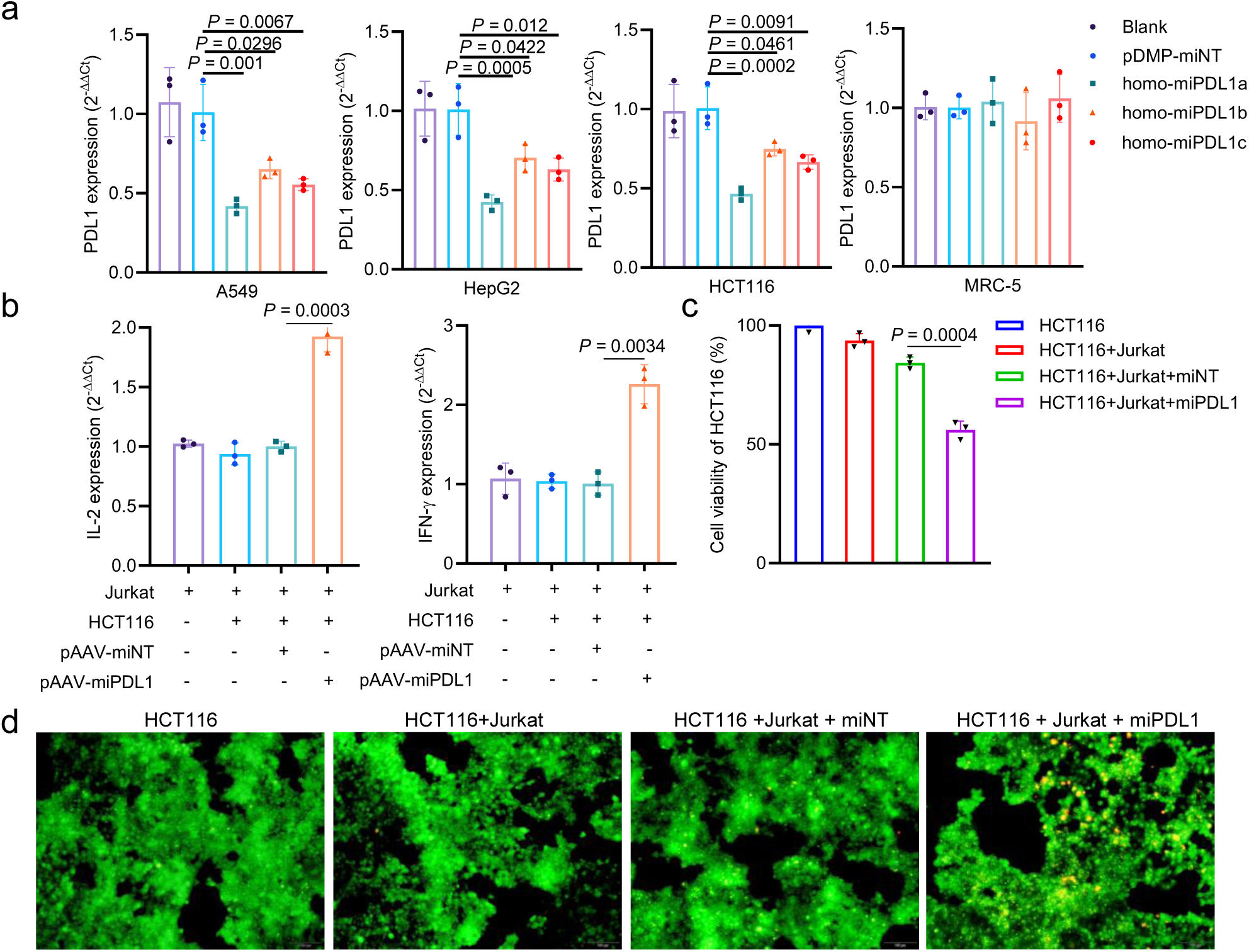
Treatment of human cells with pAAV-miPDL1 that can expressing miRs targeting human PD-L1 mRNA. **a** Human PD-L1 expression in human cells (A549, HepG2, HCT116, and MRC-5) transfected with pAAV-miPDL1s (homo-miPDL1-a, homo-miPDL1-b, and homo-miPDL1-c) detected by qPCR (n=3 wells). **b** Expression of IL-2 and IFN-γ in Jurkat cells detected by qPCR (n=3 wells). **c** Cell viability of HCT116 detected by CCK-8 reagent (n=3 wells). **d** AO&EB staining of HCT116 cells with different treatment. HCT116 cells were transfected with pAAV-miNT or pAAV-miPDL1 and then co-cultured with Jurkat (**b**, **c**, and **d**).

**Fig. S8.**
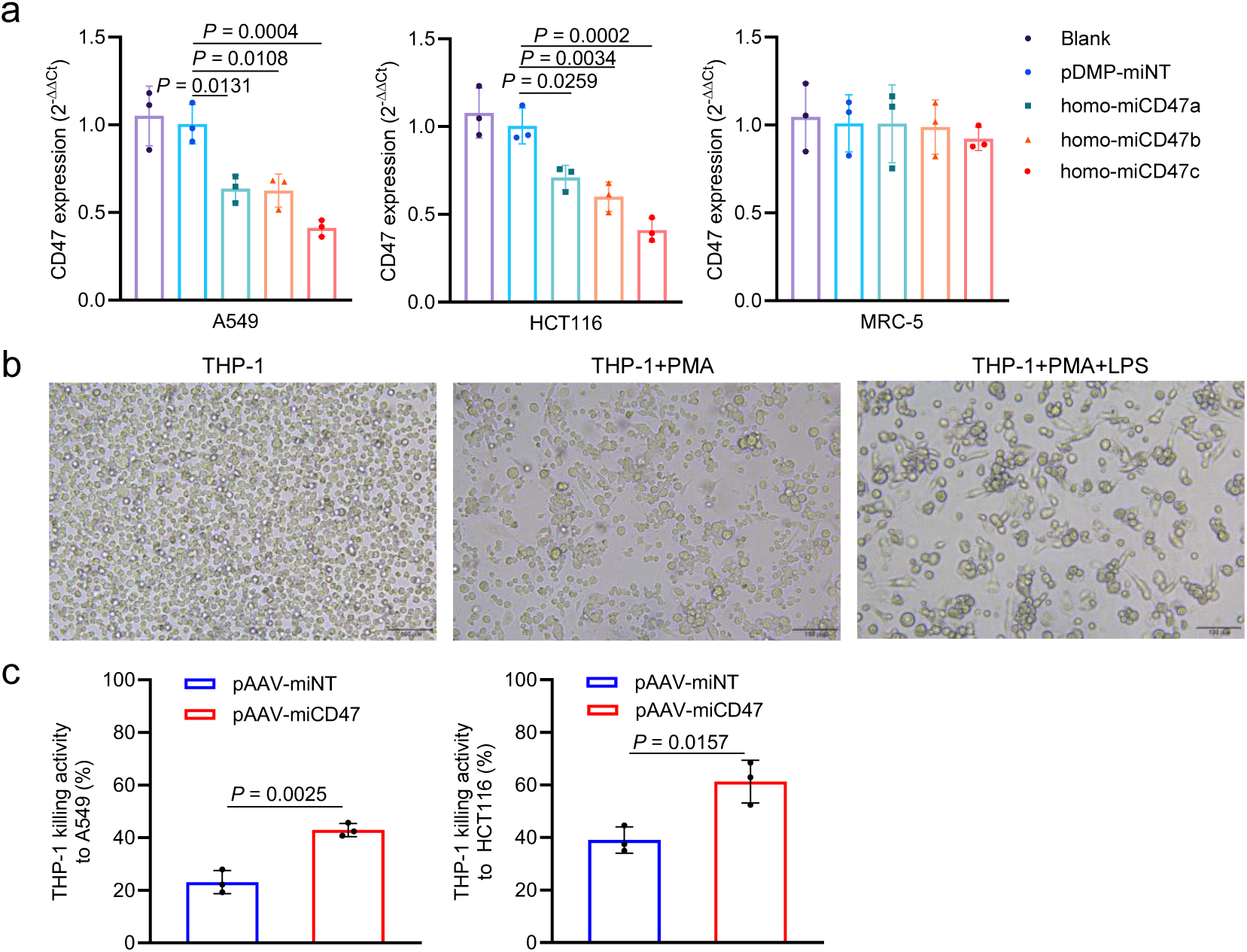
Treatment of human cells with pAAV-miCD47 that can expressing miRs targeting human CD47 mRNA. **a** Human PD-L1 expression in human cells (MRC-5, A549, and HCT116) transfected with pAAV-miCD47 (homo-miCD47-a, homo-miCD47-b, and homo-miCD47-c) detected by qPCR (n=3 wells). **b** THP-1 cell culture and graded activation with PMA and LPS. **c** Cell viability of A549 and HCT116 cells co-cultured with THP-1 detected by CCK-8 reagent (n=3 wells). Cells were transfected with pAAV-miNT or pAAV-miCD47 and then co-cultured with the activate THP-1 cells.

**Fig. S9.**
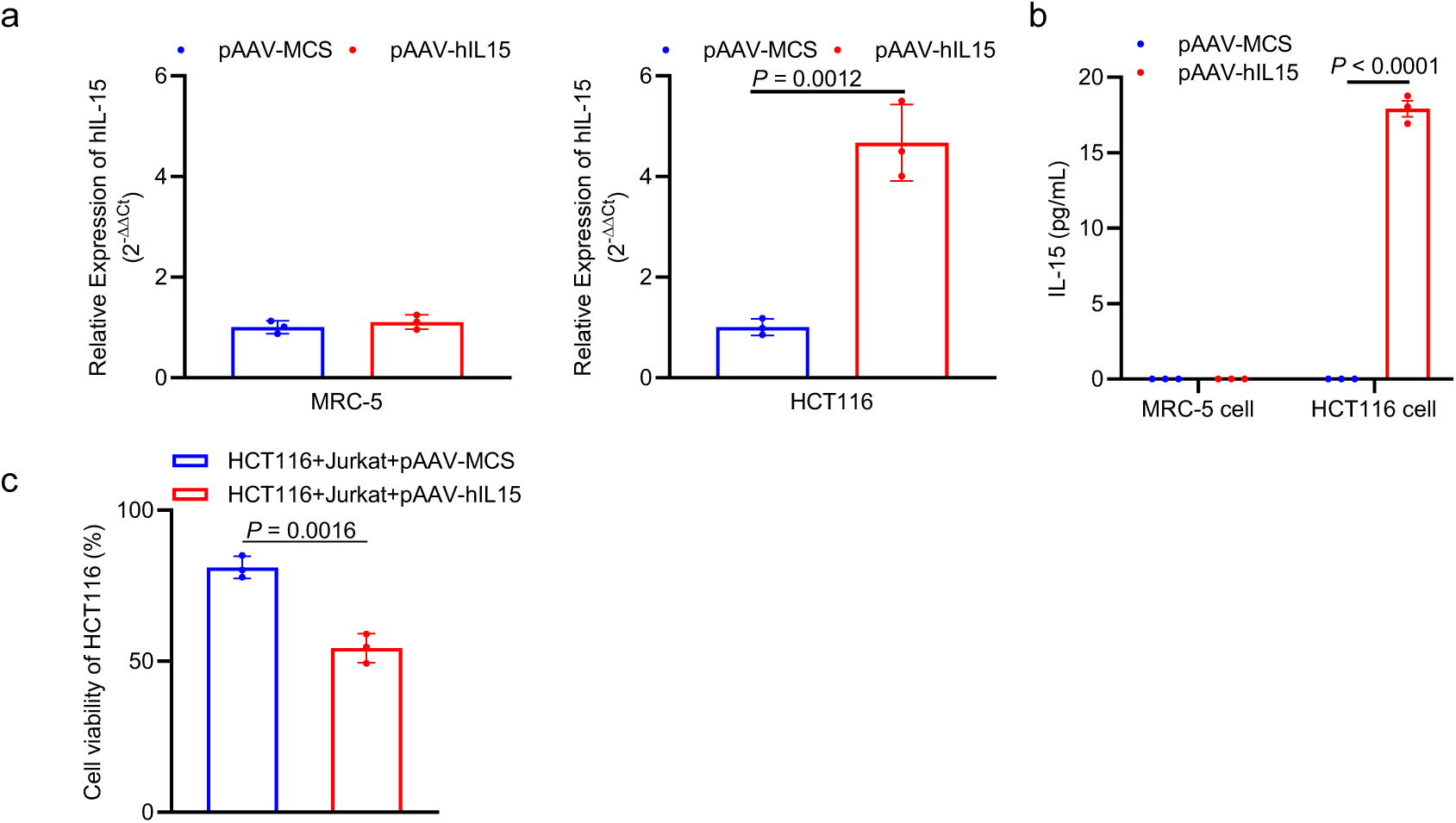
Treatment of human cells with pAAV-hIL15. **a** Human IL-15 expression in human cells (MRC-5 and HCT116) transfected with pAAV-hIL15 detected by qPCR (n=3 wells). **b** IL-15 expression in MRC-5 cells and HCT116 cells detected by ELISA (n = 3 wells). **c** Cell viability of HCT116 with Jurkat cells co-culture, detected by CCK-8 reagent (n=3 wells).

## Notes

### Competing Interest Statement

The authors have declared no competing interest.

